# Stochastic synchronization of dynamics on the human connectome

**DOI:** 10.1101/2020.02.09.940817

**Authors:** James C. Pang, Leonardo L. Gollo, James A. Roberts

## Abstract

Synchronization is a collective mechanism by which oscillatory networks achieve their functions. Factors driving synchronization include the network’s topological and dynamical properties. However, how these factors drive the emergence of synchronization in the presence of potentially disruptive external inputs like stochastic perturbations is not well understood, particularly for real-world systems such as the human brain. Here, we aim to systematically address this problem using a large-scale model of the human brain network (i.e., the human connectome). The results show that the model can produce complex synchronization patterns transitioning between incoherent and coherent states. When nodes in the network are coupled at some critical strength, a counterintuitive phenomenon emerges where the addition of noise increases the synchronization of global and local dynamics, with structural hub nodes benefiting the most. This stochastic synchronization effect is found to be driven by the intrinsic hierarchy of neural timescales of the brain and the heterogeneous complex topology of the connectome. Moreover, the effect coincides with clustering of node phases and node frequencies and strengthening of the functional connectivity of some of the connectome’s subnetworks. Overall, the work provides broad theoretical insights into the emergence and mechanisms of stochastic synchronization, highlighting its putative contribution in achieving network integration underpinning brain function.

## 1. Introduction

Oscillatory networks are ubiquitous in nature and are the core of several biological systems such as gene-regulatory networks and brain circuits [1, 2]. Their overarching functions rely on how the collective dynamics of their constituent nodes (often measured in terms of synchronization) self-organize due to mutual interactions and/or from entrainment to external signals. For example, modern power grids can operate normally only if all generators of the network are stably synchronized at the same frequency [3]. Similarly, neural synchronization is central to many aspects of brain function, including attention and information transfer [4, 5, 6].

Network synchronization crucially depends on several factors such as network architecture, intrinsic frequency of the oscillators, strength of the interaction between oscillators, and external inputs. These factors have been studied in models; perhaps the most famous is the Kuramoto model [7], which considers the phase synchronization of coupled oscillatory units. The simplest Kuramoto model, which assumes a fully connected network, can produce nontrivial transitions into or out of synchronization [8], analogous to phase transitions studied in statistical physics [9]. Several theoretical works have then leveraged the analytical tractability of the model to exactly solve synchronization states in small artificial systems [10] and infinitely-sized systems with a symmetric distribution of oscillator frequencies [11, 8]. Since then, the model has been extended to more realistic systems such as networks with heterogeneous connectivity, e.g., small-world and random networks [12, 13], networks with delay-dependent couplings [14], networks with time-delayed interactions [15, 16, 17], and networks influenced by extrinsic structured or stochastic inputs [18, 19].

In this work, we study synchronization in the human brain, which is considered to be a large interconnected complex network that operates at multiple scales in space and time [20]. Macroscopically, the brain comprises anatomical white matter connections between cortical regions called the connectome [21, 22], serving as the substrate that shapes largescale neuronal dynamics. These connections are heterogeneous in nature such that certain brain regions possess relatively larger number of connections (termed hubs) [23], which have been shown to facilitate integration of activity [24]. In addition, neuronal dynamics exhibit heterogeneous timescales [25], such that higher-order brain hubs exhibit slower oscillations [26]. This structure–dynamics relationship in the brain drives the generation of local and global network synchronization patterns that support the brain’s vital functions [27, 28]. Indeed, the use of coupled oscillators to generate large-scale brain dynamics has reproduced many observed features in neuroimaging data [29, 30, 31, 32, 33, 34]. Nevertheless, the complex links between brain structure and function remain incompletely understood. This is important to address not just for understanding the workings of normal brain function but also those of pathologies, such as epilepsy [35] and autism [36], which are linked to excess or deficits in cortical synchronization.

Network synchronization is also inevitably influenced by noise or stochastic perturbations. Noise is intuitively considered to be detrimental to overall synchronization. However, paradoxical effects exist in various physical and biological systems, where a moderate amount of noise can affect the system’s dynamics by inducing or enhancing synchronization. This is known as noise-induced synchronization or stochastic synchronization [37, 38]; note that both terms interchangeably used in the literature, and we elect to use the latter in this study. Such roles for noise bear similarity to the well-known stochastic resonance in physics, where noise can optimize the response of nonlinear systems to a weak external input [39]. Noise can also aid the synchronization of periodic [40, 41] and chaotic [42] oscillators and networks of oscillators with various topologies [43, 44]. Noise can also increase the regularity of activity in excitable systems [45]. Finally, in neural systems, noise can induce phase-entrainment of neural waves [46], synchronize uncoupled neural oscillators [47], improve sensorimotor performance [48], and enhance the firing synchronization of Hodgkin-Huxley neurons [49]. Even with this evidence of the presence and benefits of noise in physical and biological systems, stochastic synchronization and its mechanisms have rarely been investigated in the context of large-scale biological networks, especially in the human connectome, and hence is likely of interest for the neuroimaging community.

In this work, we aim to systematically understand the effect of stochastic perturbations to oscillators on the human connectome that follow a hierarchy of timescales. That is, the model incorporates heterogeneous, hierarchically distributed oscillatory frequencies and a complex connectivity (weight distribution), such that highly connected hub regions have slow natural frequencies and peripheral (nonhub) regions have fast natural frequencies [50]. Moreover, we extensively analyze and tease apart the contributions of the connectome’s topological structure and intrinsic dynamics to the emergence of stochastic synchronization, elucidating its general mechanisms.

## 2. Results

### 2.1. Brain network model

We model whole-brain oscillations using a large-scale network of coupled Kuramoto oscillators [7, 8, 51]. The phase dynamics *θ_j_* of oscillator *j* are governed by the stochastic differential equation

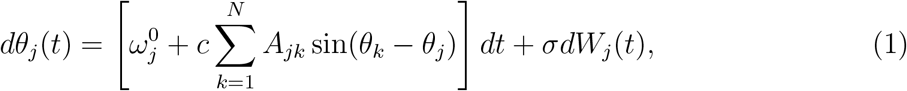

where *N* is the total number of oscillators. The model has three main ingredients: (i) connectivity matrix *A* where element *A_jk_* describes the strength of the connection from oscillator *k* to *j*, (ii) natural frequency of oscillation 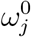, and (iii) noise represented by the Wiener process *dW_j_*. In addition, the model has two parameters: (i) coupling strength *c* that scales the connection strengths between oscillators and (ii) noise strength *σ* that scales the noise amplitude. Here, all nodes in the network are perturbed by independent Gaussian noise realizations with the same standard deviation *σ*.

For the connectivity matrix *A*, we use a human structural connectome derived from healthy participants [52, 53]. It is a weighted and symmetric matrix (Fig. 1A), with each row or column representing a brain region that aggregates populations of neurons, and each connection *A_jk_* is related to the number of fibers reconstructed between region *j* and *k* (see Materials and Methods for details). It has several features such as a hierarchical-modular organization [22, 54] (see solid boxes in Fig. 1A) and a spatial embedding characterized by an exponentially decaying edge weight-fiber length relationship [52, 55].

**Figure 1:**
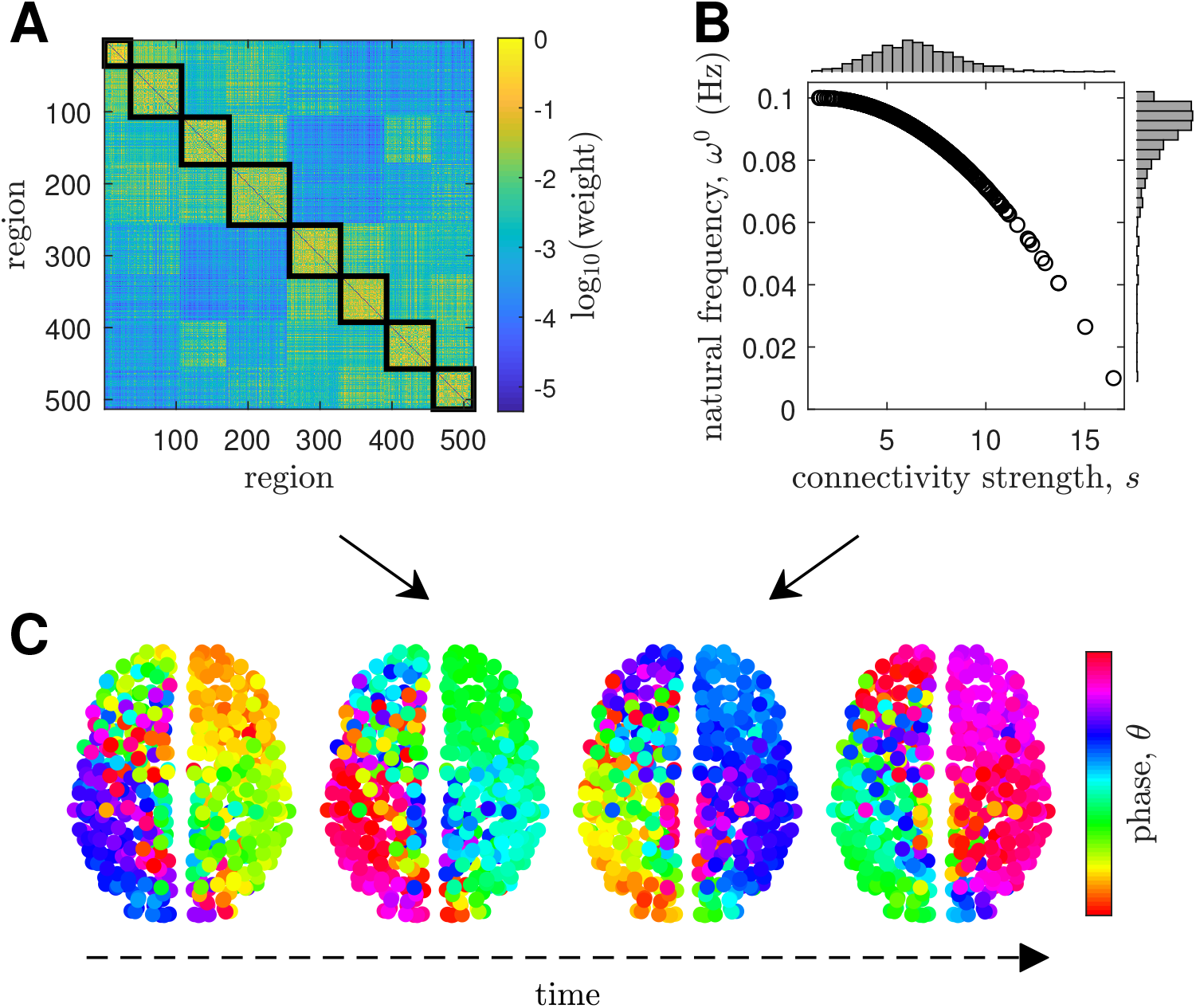
Whole-brain network model. **(A)** Connectivity matrix. The solid boxes denote modules obtained via a community detection algorithm. **(B)** Natural frequency of oscillation *ω*^0^ as a function of connectivity strength *s*. The distributions of *s* and *ω*^0^ are shown above and to the right of the panel, respectively. **(C)** Sample spatial distribution of node phases *θ* through time using the model ingredients in panels A and B.

The natural frequencies *ω*^0^ are drawn from a distribution *g*(*ω*^0^), typically chosen as a symmetric function, such as a Gaussian or Lorentzian distribution, to obtain analytically tractable solutions [8]. However, the novelty of our study is a more complex but also more neurobiologically motivated choice of *g*(*ω*^0^). We use an asymmetric hierarchical distribution, which is inspired by recent studies showing that cortical areas of primates have intrinsic timescales that differentiate according to the anatomical hierarchy [25, 56]. We determine the natural freαuency 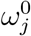 of node *j* as [57, 33]

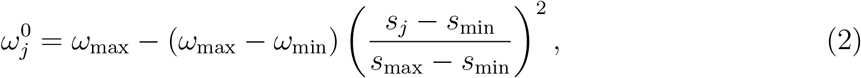

where *s_j_* is the node’s connectivity strength given by *s_j_* ∑_*k*_ *A_jk_*, frequency limits *ω*_min_ = 0.01 Hz and *ω*_max_ = 0.1 Hz are set to match the frequency bandwidth of fMRI, *s*_min_ is the minimum of all node strengths, and s_max_ is the maximum of all node strengths. This distribution, which showed the best fit to empirical resting-state fMRI data [57], allows the cortical hierarchy to be mapped into a gradient of timescales with slow hub and fast nonhub regions. The resulting 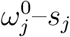 relationship for all nodes in the connectome is shown in Fig. 1B.

The above two model ingredients provide the structure and dynamics of the human brain, which when combined allows phase oscillations of nodes in the connectome to evolve through mutual interactions without the need for any external input (Fig. 1C). We then thoroughly explore the effect of coupling strength and noise on these interactions.

Note that our formulation in Eq. (1) does not include the effects of time-delayed neural interactions, which are often considered in whole-brain network models to capture delays in signal transmission between brain regions due to long white matter tracts [16, 17, 26, 34]. However, since time delays in the brain are approximately in the order of 10 ms [17, 34] and we focus on fMRI dynamics with slow timescales (periods of oscillations in the range of 10 to 100 s), neglecting time delays would not significantly affect the dynamical solutions of Eq. (1) and the overall network oscillations.

### 2.2. Network state transitions

We first characterize the effect of the coupling strength *c* and the noise strength *σ* on the collective network dynamics measured in terms of three quantities: coherence *R*, synchronization *S*, and metastability *M*. Briefly, coherence *R* captures the phase alignment of all oscillators in the network at each instance in time, with its value varying between 0 (fully incoherent state) and 1 (fully coherent state). Synchronization *S* and metastability *M* are summary statistics at steady state capturing the mean and variability of *R*, respectively (see Materials and Methods for details).

To understand the fundamental behavior of the network without any external influences, we first neglect noise (i.e., *σ* = 0). Figure 2A shows the network’s *S* and *M* for varying coupling strengths. Our results reveal that nodes in the network organize into diverse states of collective dynamics by just tuning *c*. In particular, weak coupling (*c* ≤ 0.001) consistently leads to incoherent states with low *S*, while stronger coupling (*c* ≥ 0.006) leads to coherent states with high S. More importantly, we find a regime placed between the incoherent and coherent states where partial synchronization emerges. Coherence dynamics in this intermediate regime have high-amplitude fluctuations, demonstrated by high *M* with peak occurring at a critical value *c*\* = 0.0027. These fluctuations are clearly demonstrated by the time evolution of *R* and local spatiotemporal phase dynamics at *c* = *c*\* = 0.0027 shown in Fig. 2B and Fig. 2C, respectively (see Fig. S1 to compare *R* and phase dynamics at *c* ≪ *c*\*, *c* = *c*\*, and *c* ≫ *c*\*). This critical coupling strength places the network in a flexible regime that spontaneously accesses both states of order and disorder. Moreover, this critical coupling strength is very close to the value (i.e., *c* = 0.003) obtained when the same network model is fitted to resting-sate fMRI data [57, 33]. Hereafter, unless otherwise stated, we fix the coupling strength to *c* = *c*\* = 0.0027 to further investigate network dynamics on the human connectome at this critical point.

**Figure 2:**
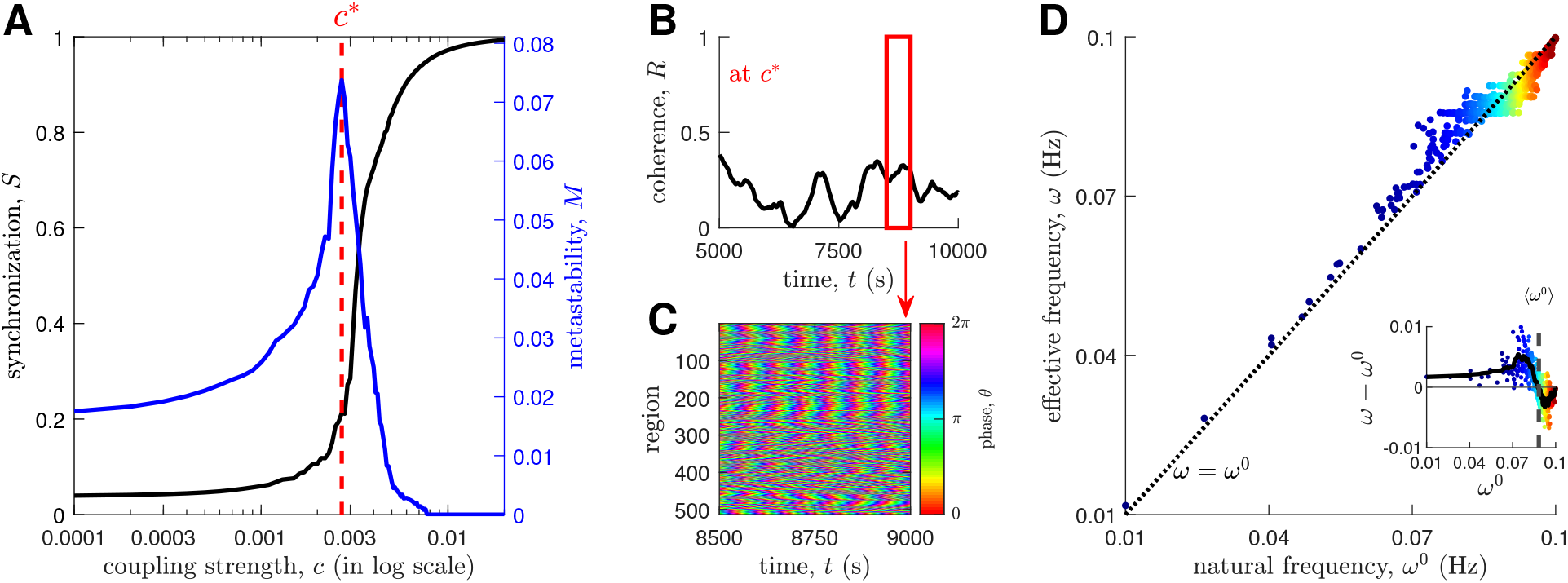
Noise-free network dynamics. **(A)** Network synchronization *S* and metastability *M* vs coupling strength *c*. The red dashed line highlights the critical coupling strength *c\** where *M* is maximum. The solid lines represent ensemble averages of 30 initial conditions. **(B)** Time evolution of network coherence *R* at *c*\*. **(C)** Local phase dynamics within the time window highlighted by the red solid box in panel B. **(D)** Effective node frequency *ω* as a function of natural frequency *ω*^0^ at zero noise. The markers are colored according to the order of natural frequencies (blue: low *ω*^0^; red: high *ω*^0^). The dashed line represents *ω* = *ω*^0^. **(Inset)** *ω* — *ω*^0^ as a function of *ω*^0^. The solid line represents the sliding-window mean smoothed by a Savitzky-Golay filter (order 3 and length 15) and the dashed line represents the mean-ensemble frequency 〈*ω*^0^〉.

To further analyze the effect of placing the network at the critical point, we calculate the effective node frequency *ω* (see Materials and Methods for details) [58]. Figure 2D shows that due to node interactions even without any external influences, nodes in the network adjust their frequencies and the strength-frequency relationship imposed in Fig. 1B becomes weaker. In fact, nodes originally oscillating below/above the mean-ensemble frequency [59] of 〈*ω*^0^〉 = 0.0883 Hz tend to speed up/slow down (see inset of Fig. 2D). That is, hubs speed up and peripheral (nonhub) regions slow down. Moreover, Fig. 2D clearly shows the formation of frequency clusters, demonstrating that interactions alone force some nodes to align their frequencies of oscillation (see formation of horizontal lines near the mean-ensemble frequency).

### 2.3. Emergence of stochastic synchronization

The network responses described above demonstrated the ability of the model to generate complex patterns of collective dynamics across the human connectome by just tuning the coupling strength. The dynamics were achieved only via mutual interactions without the need for external inputs. Hence, the next question we want to address is how the network would respond to inputs. More specifically, how would stochastic perturbations (i.e., independent additive noise at each node) affect network dynamics and synchronization? We systematically answer this question by characterizing the effects of changes in the noise strength σ.

Initially, we analyze the network’s behavior for three noise strengths: the original case without noise (*σ* = 0; Supplementary Movie S1); moderate noise (*σ* = 0.008; Supplementary Movie S2); and high noise (*σ* = 0.2; Supplementary Movie S3). On a node level, visual inspection of the spatial distribution of node phases (Fig. 3A) reveals that larger clusters of similar node phases are more apparent for the moderate noise strength. This is quantitatively supported by a narrower distribution of phase differences Δ*θ* centered at Δ*θ* = 0 (see Fig. S2B). On a network level, we find that the network’s coherence paradoxically increases when the noise strength is raised to a moderate value compared to the noise-free case, but breaks down for a higher value (Fig. 3B). This phenomenon is further demonstrated by the probability density function (pdf) and the mean of *R* (Figs 3C and 3D), showing concentrations at higher R values for the moderate noise strength (with statistically significant pairwise differences between the (R) of the three noise levels via one-way ANOVA and multiple comparison testing; *p* < 1 × 10^-9^).

**Figure 3:**
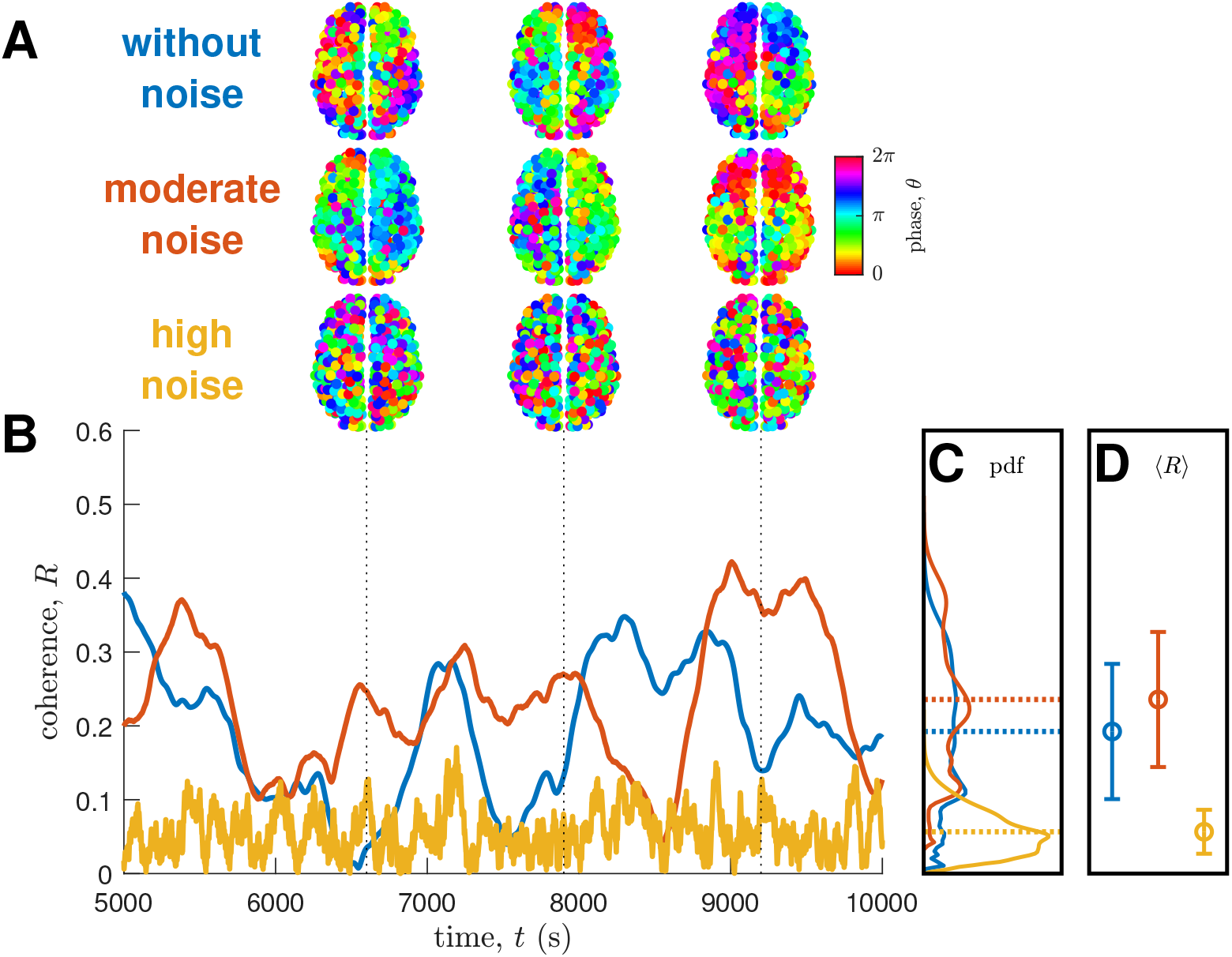
Node phases and network coherence for various noise strengths. **(A)** Spatial distribution of node phases *θ* at different time points in superior axial brain view. **(B)** Time evolution of network coherence *R.* **(C)** Kernel density estimate of the probability density function (pdf) of *R*. The dotted lines represent the mean values of *R*. **(D)** Mean and standard deviation of the pdfs in panel C. For panels B, C, and D, the lines and markers are colored according to noise strength (blue: without noise; red: moderate noise; yellow: high noise).

To further investigate the above phenomenon, we systematically vary the noise strength and calculate the percent change in network synchronization ΔS, which measures changes in synchronization with respect to the baseline noise-free case. Results in Fig. 4A show that there exists a range of noise strengths (0 < *σ* < 0.033; this includes the moderate-noise regime in Fig. 3) where network synchronization is above the baseline (i.e., Δ*S* > 0). This is also true even for very low noise strengths (see inset of Fig. 4A). Further increases in noise strength (this includes the high-noise regime in Fig. 3) lead to decreased synchronization (i.e., Δ*S* < 0), in line with the intuition that strong noise disrupts synchrony. Crucially, the phenomenon only exists when the network is placed around the critical coupling strength *c*\* (see Fig. S3). We further verify that the phenomenon is robustly found on individual human connectomes from an independent dataset from the Human Connectome Project (HCP; see Materials and Methods for details) [53] and also for different numbers of nodes in the parcellation used to construct the connectomes (see Figs S4–S6).

**Figure 4:**
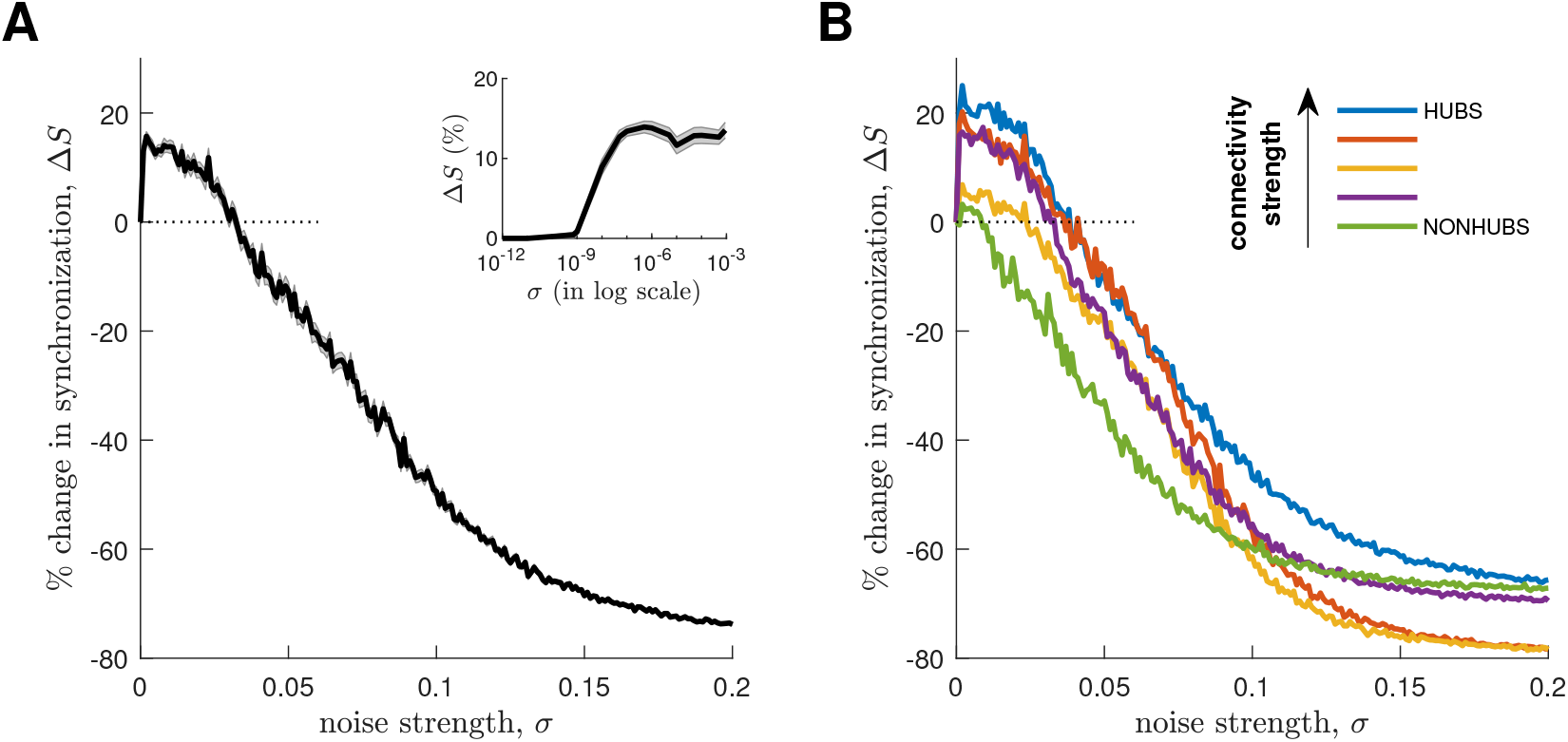
Stochastic synchronization. **(A)** Percent change of global network synchronization Δ*S* vs noise strength *σ*. **(Inset)** Δ*S* for very low noise strengths (*σ* < 1 × 10^-3^). For panel A and its inset, the solid line represents an ensemble average of 50 noise realizations and the shaded area represents the standard error of the mean. **(B)** Δ*S* vs *σ* for different subnetworks. The subnetworks are derived by partitioning the connectome into five groups according to connectivity strength s, where the top 20% represents strongly connected hub nodes and the bottom 20% represents weakly connected nonhub nodes. The solid lines represent ensemble averages of 50 noise realizations.

We next want to understand whether stochastic synchronization uniformly occurs in the whole connectome network or whether its manifestation varies locally within smaller sub-networks across the connectome’s topological hierarchy (i.e., are some subnetworks more or less synchronized by noise?). Hence, we subdivide the nodes into five groups according to the connectivity strength *s*, such that the first group (top 20%) represents hubs (nodes with high-strength connections) and the last group (bottom 20%) represents nonhubs (nodes with low-strength connections), and analyze their synchronization patterns. We find that stochastic synchronization exists heterogeneously across the connectome’s local subnetworks (Fig. 4B). In particular, hubs have higher maximum Δ*S* and a wider range of noise strengths producing Δ*S* > 0 as compared to nonhubs. This demonstrates that stochastic synchronization emerges on the connectome both globally and locally (and heterogeneously). More importantly, these results indicate that stochastic synchronization may be a useful mechanism for improving overall network integration facilitated by different subnetworks of the connectome.

### 2.4. Drivers of stochastic synchronization

In the above, we showed that noise can increase the synchronization of oscillations on the human connectome. We dig deeper and investigate the drivers of this phenomenon. Going back to the ingredients of the model, we conjecture that the paradoxical phenomenon could be due to the intrinsic node frequencies and/or the rich network topology of the human connectome. Thus, we explore two scenarios.

The first scenario is using the human connectome but changing the distribution of natural frequencies to other well-known distributions typically used in the Kuramoto model literature: (i) Dirac-delta (homogeneous), (ii) random uniform (rand-uniform), (iii) random Gaussian (rand-gaussian), and (iv) random Lorentzian (rand-lorentzian). We constrain them to the same frequency bandwidth as in Fig. 1B (their pdfs are shown in Fig. 5A; see Materials and Methods for further details). Unlike the hierarchical case, these distributions have no consistent relationship between the network’s topology and the natural frequencies. We set the coupling strength to *c* = 0.0027, which is the *c*\* found in Fig. 4, and determine changes in the coherence patterns in the absence of noise (Fig. 5B). Most notably, the homogeneous distribution leads to highly coherent dynamics (as it must [8]) and the rand-lorentzian distribution leads to dynamics that are similar to the original hierarchical case. However, stochastic synchronization does not occur for the other distributions, in fact desynchronization is observed with addition of noise (Fig. 5C). We verify that the results hold even when the number of realizations used in Fig. 5C is increased (see Fig. S7). This reveals that it is important for the human connectome to have its nodes oscillate according to their anatomical hierarchy in order for stochastic synchronization to emerge.

**Figure 5:**
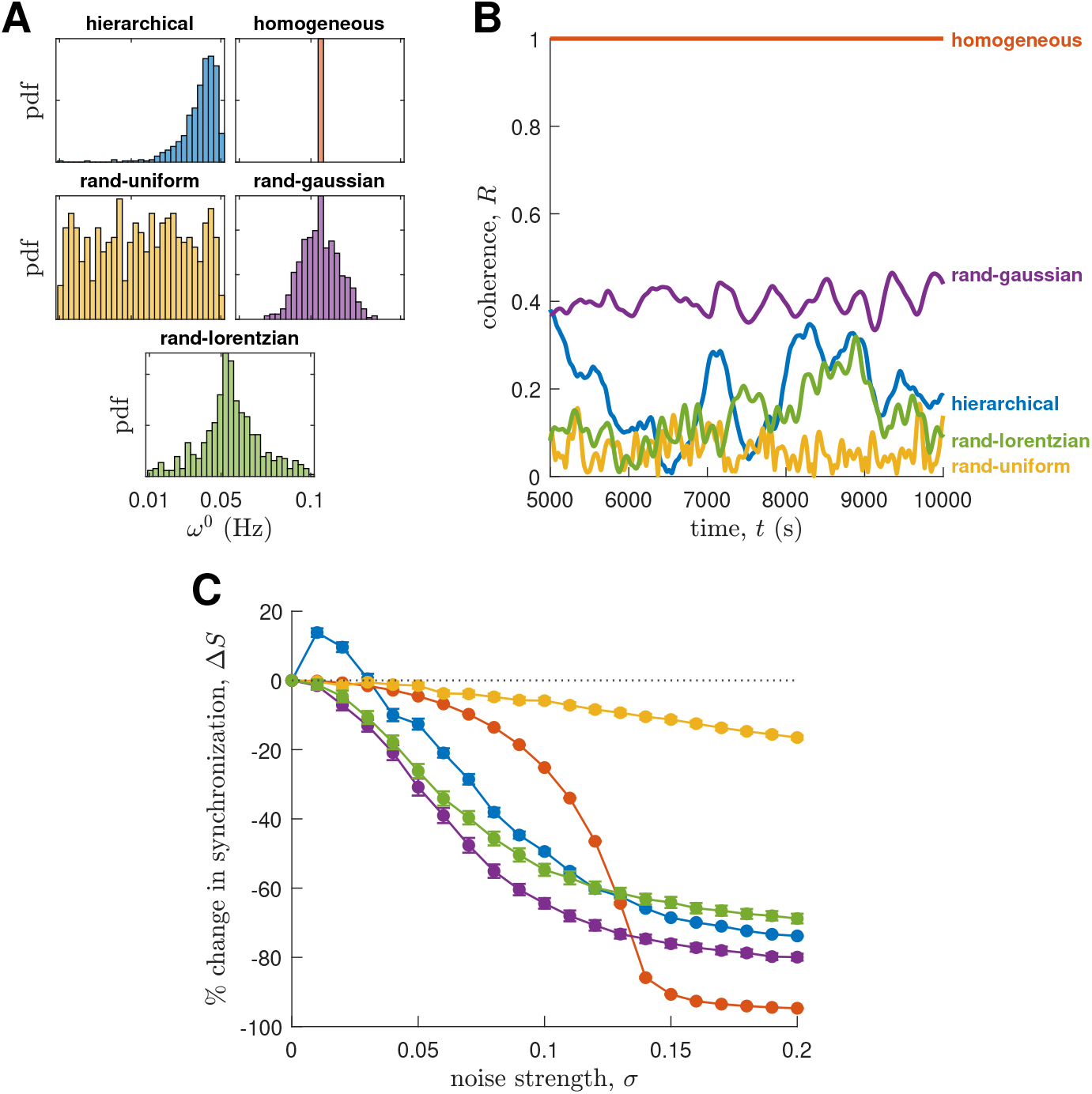
Role of the distribution of natural frequencies on synchronization dynamics. **(A)** Probability density function (pdf) of different frequency distributions. **(B)** Time evolution of network coherence R without noise. The lines are colored as labeled and according to panel A. **(C)** Percent change of network synchronization Δ*S* vs noise strength *σ*. The lines are colored according to panel A. The markers represent ensemble averages of 50 realizations of the frequency distributions and the vertical lines represent the standard errors of the means.

We note that thus far we have analyzed the synchronization of network dynamics at the coupling strength value of *c* = 0.0027, which is the *c*\* specific to the human connectome-hierarchical distribution pair (obtained from the peak *M* in Fig. 2A). However, it is possible that the value of *c*\* changes specific to the network-frequency distribution pair used. Therefore, we ask whether the stochastic synchronization effect would emerge in Fig. 5C if we properly tune the different human connectome-frequency distribution pairs to their respective *c*\*. Hence, we recalculate the *S* and *M* vs *c* curves (similar to Fig. 2A) specific to the different frequency distributions and measure *c*\* corresponding to the peak *M* (the results are shown in Fig. S8A). Then, we simulate the network dynamics using the new *c*\* and recalculate the synchronization vs noise strength curves (the results are shown in Fig. S8B). By doing so, the results in Fig. 5C generally still hold, albeit the rand-lorentzian distribution weakly showing the effect with maximum ΔS of ≈1%. These results highlight the robustness of our findings and that the hierarchical frequency distribution is truly an important driver to strongly produce the effect. Moreover, these results further show that the effect only exists when the oscillations of a complex network are governed by a complex frequency distribution near criticality, possibly with some sensitivity to the precise tuning of the coupling. Note that we did not perform this analysis on the homogeneous distribution case because it always leads to full network synchronization at steady state; hence, its *S* vs *c* curve is flat at *S* = 1 without its own critical coupling strength. However, we found some interesting transient dynamics for this case where noise can hasten the transition to synchrony, similar to the findings of [12] (this will be explored in a future study).

The second scenario is retaining the hierarchical frequency distribution but changing the topology of the network. We explore five alternative connectome topologies. They include traditional surrogate networks [60] as well as recently developed surrogates that respect (or partially respect) the spatial embedding of the human brain, enabling tests of the extent to which network phenomena depend on weight-distance relationships and the positions of hubs [52, 55]. The tested topologies are: (i) fully connected network; (ii) weight-preserving random network, randomizing 50% of the connections; (iii) weight-preserving random network, randomizing 100% of the connections; (iv) geometry-preserving random network, preserving the node strengths; and (v) geometry-preserving random network, preserving the node-strength sequence. Their edge weight-fiber length relationships are shown in Fig. 6A (see Materials and Methods for further details). Topology (i) is used by the simplest Kuramoto model in the literature, topologies (ii) and (iii) are surrogates commonly used in network science, and topologies (iv) and (v) are more biologically realistic surrogates [52, 55]. As above for the different frequency distributions, we set the coupling strength to *c* = 0.0027. We find that changing the network’s topology leads to various coherence patterns in the absence of noise (Fig. 6B). Most notably, the fully connected and fully randomized topologies (i and iii) lead to highly coherent dynamics, while the topology with preserved node-strength sequence (v) leads to dynamics that are similar to the human connectome. However, interestingly, we find that only the human connectome yields the stochastic synchronization effect (Fig. 6C). Despite the geometry- and strength-sequence-preserving surrogate network (light blue) having the same weight-distance relationship and hub locations as the original brain network, and exhibiting similar coherence dynamics, noise does not enhance its synchronization, implying that more subtle topological features of the brain’s wiring contribute to the effect such as possibly the brain’s small-world topological structure (see Fig. S9 and Materials and Methods for details) [43]. These results reveal that the heterogeneity in the connectivity pattern, the hierarchy of the connectivity strengths of each node (Fig. 1B), and, more importantly, the topology of the human connectome are crucial for driving the emergence of stochastic synchronization. The same conclusions remain even if we tune the different network-hierarchical frequency pairs to their respective critical coupling strengths (see Fig. S10).

**Figure 6:**
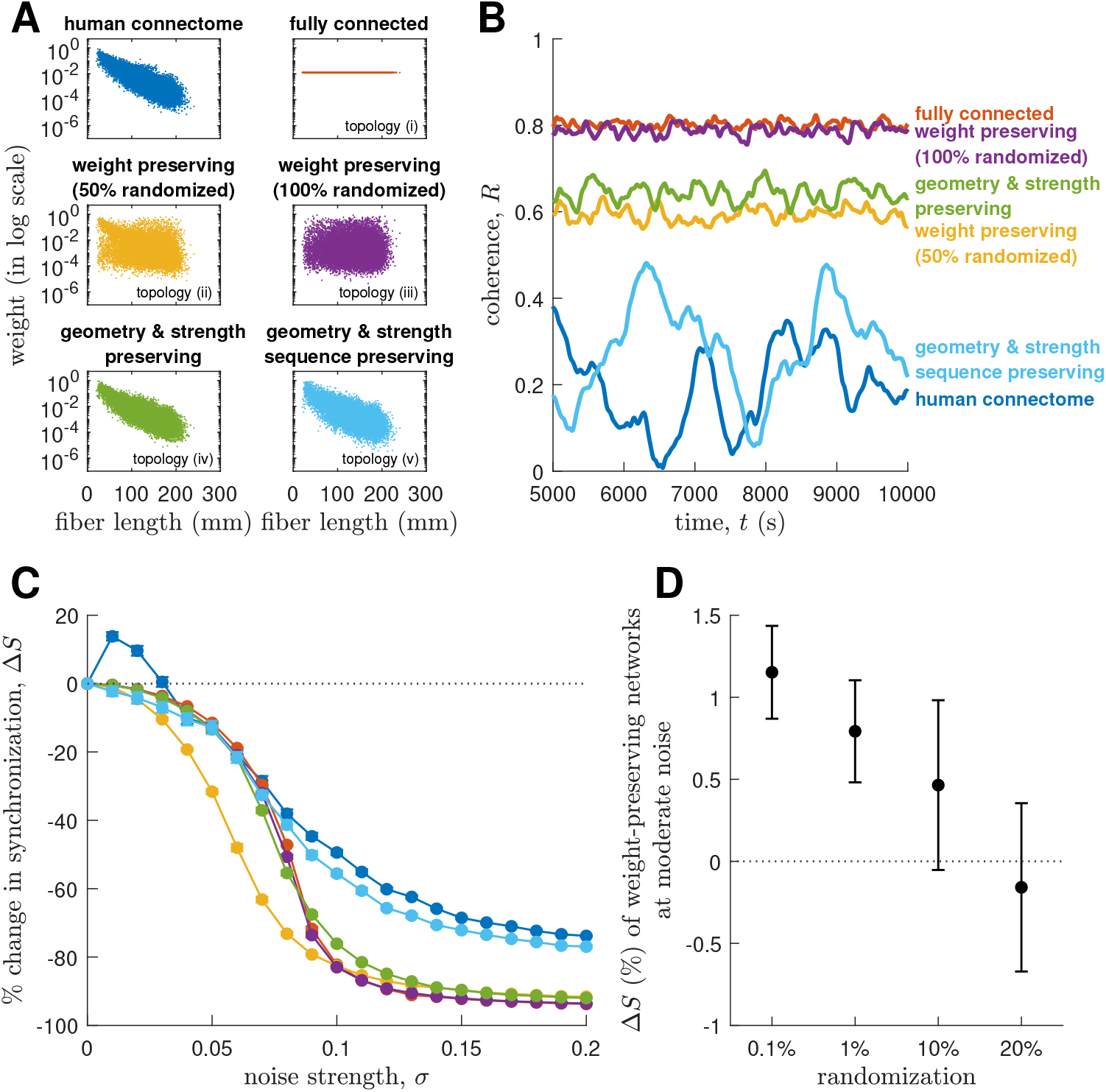
Role of the network’s topology on synchronization dynamics. **(A)** Log of edge weight vs fiber length for all pairs of regions of different network topologies. **(B)** Time evolution of network coherence R without noise. The lines are colored as labeled and according to panel A. **(C)** Percent change of network synchronization Δ*S* vs noise strength *σ*. The lines are colored according to panel A. The markers represent ensemble averages of 50 surrogates of the network topologies and the vertical lines represent the standard errors of the means. **(D)** Percent change of network synchronization Δ*S* of different weight-preserving networks (with a small to moderate fraction of randomized weights) at moderate noise. The markers represent ensemble averages of 500 surrogates of the network topologies and the vertical lines represent the standard errors of the means.

We next sought to determine how far from the empirical network the stochastic synchronization effect persists by rewiring a subset of edges. In particular, we generate weight-preserving network surrogates of the human connectome [similar to topologies (ii) and (iii)] but only randomize a small fraction of edges than before [52, 55], that is, a range of randomizations of ≤20%, to delineate the robustness of the effect with distance from the human connectome’s topology. Figure 6D shows that the effect can be robustly produced when we randomize a small to moderate fraction of weights (0.1%, 1%, and 10%) albeit it is not as strong as what we have found for the human connectome in Fig. 4A where the maximum Δ*S* was ≈16%. Then, as we increase the randomization to 20% [and beyond, e.g., topologies (ii) and (iii)], the effect disappears and Δ*S* becomes negative. These results demonstrate that it is possible to produce the effect using other network topologies, especially networks that closely resemble the human connectome. In addition, we find that these networks that also produce the effect have similar levels of “small-worldness” as the human connectome (Fig. S9), which could be an important topological property shared by these networks that is informative of the emergence of the effect.

### 2.5. Mechanisms of stochastic synchronization

At this point, we have explored the role of the connectome’s structure and the nodes’ intrinsic frequencies on the emergence of stochastic synchronization. Alongside these important ingredients, we ask what general mechanisms might explain the occurrence of this phenomenon. We investigate two lines of inquiry, one based on local node dynamics and the other based on large-scale network dynamics.

To investigate the phenomenon based on local node dynamics, we analyze two things: the nodes’ phases and the nodes’ effective frequencies of oscillations.

We first analyze how the nodes’ phases cluster together in line with standard phaselocking calculations for oscillators [61] and clustering analyses in neuroimaging studies [62]. We do this by placing node *j* on a unit circle at location (*x_j_, y_s_*) defined by its phase *θ_j_* at time *t* such that (*x_j_,y_j_*) = (cos*θ_j_*, sin*θ_j_*) (Fig. 7A). We perform this for all nodes in the network through time and quantify the tendency of nodes with similar phases to cluster (Fig. 7B). We investigate phase-clustering using a density-based clustering algorithm that groups together points that are close to each other within a threshold based on a distance measurement; in our case, we use the cosine distance to take into account distances on a circle (see Materials and Methods for details). We then calculate the average number of clusters formed and the average size (number of nodes) of the largest cluster across time after transients have been removed, and compare the trends of these values to the synchronization of the network for different noise strengths. In agreement with Fig. 4, the network synchronization improves for most of the noise realizations tested when the network is perturbed by noise with moderate strength and then significantly deteriorates for noise with high strength (Fig. 7C). Moreover, stochastic synchronization coincides with the entrainment of node phases to align with each other, resulting, on average, in fewer phase clusters (Fig. 7D) but with bigger sizes (Fig. 7E). Comparing the statistics of the high-noise case with those of two null models (i.e., uncoupled and random; see Materials and Methods for details), we find that a high noise strength allows noise to dominate the dynamics of the network such that synchronization ceases and the network is pushed to a configuration as if the nodes are disconnected and the phases of their oscillations are purely random. This finding for the high-noise case is further corroborated in Fig. S11, showing that the nodes return to their respective natural frequencies.

**Figure 7:**
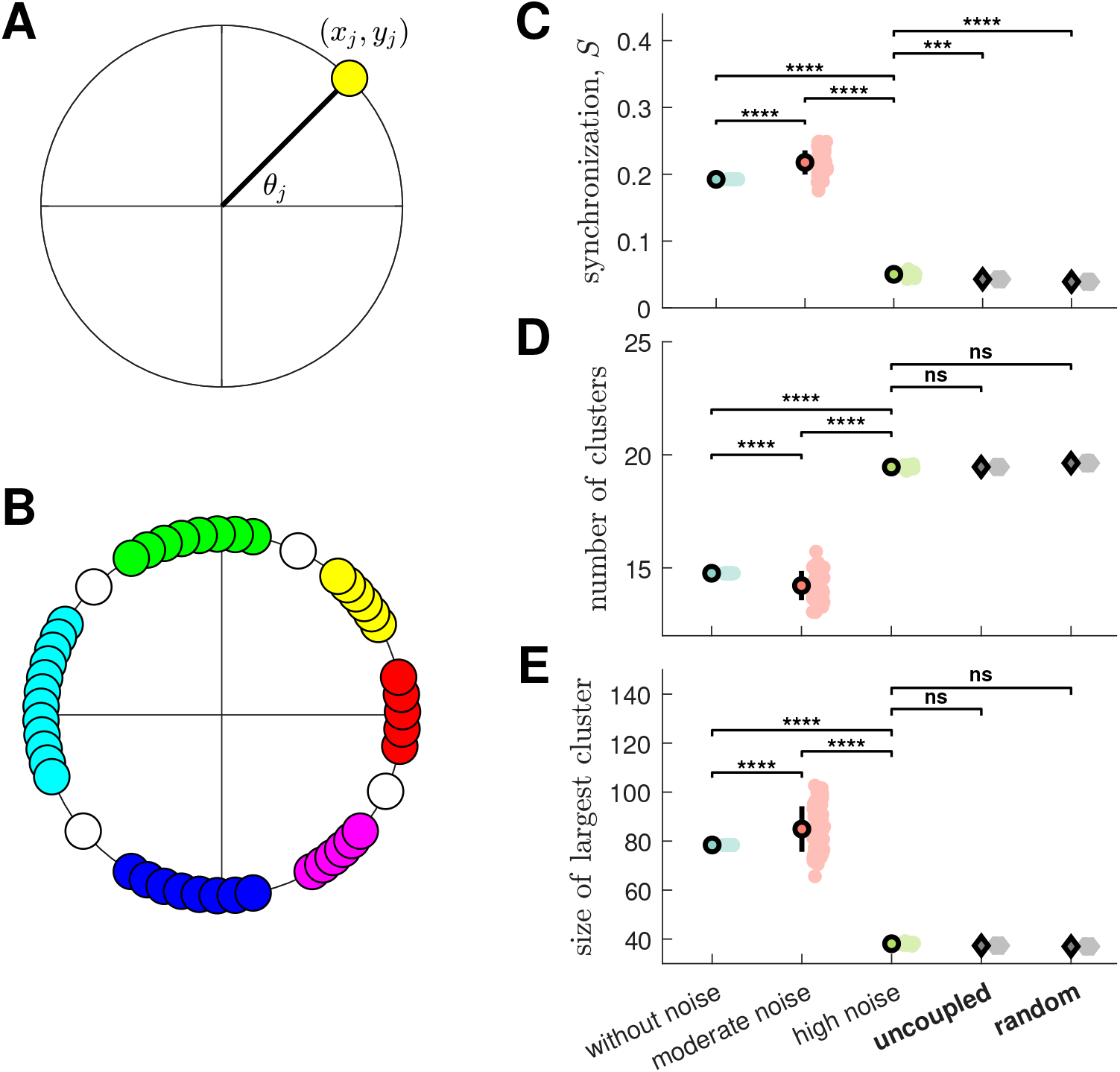
Synchronization and phase-clustering statistics for various noise strengths and null models. The two null models are uncoupled, where all nodes are disconnected and node phases evolve independently, and random, where all node phases are randomly distributed. **(A)** Schematic of a node *j* with phase *θj* at location *(xj,yj*) on a unit circle. **(B)** Schematic of phase clustering, where the nodes are colored according to the cluster they belong to. Uncolored nodes do not belong to any cluster. **(C)** Synchronization *S*. **(D)** Number of clusters. **(E)** Size of largest cluster. For panels C, D, and E, the clouds of points represent 50 noise realizations, the thick markers represent ensemble averages of all noise realizations, and the vertical lines represent standard deviations. The two null models are in bold face. **** and *** denote a statistically significant difference with *p* < 10^-4^ and *p* < 10^-3^, respectively, while ns denotes no significant difference.

Next, we analyze the nodes’ effective frequencies for the range of noise strengths investigated in Fig. 4. Figure 8A shows that, in general, noise influences nodes to change their frequencies, with the changes dependent on the strength of input noise. Moreover, the changes in frequencies are more clearly observed from *ω* = 0.08 to 0.1 Hz, where the mean-ensemble frequency 〈*ω*^0^〉 = 0.0883 Hz sits. Hence, to better understand Fig. 8A, we focus on the *ω* = 0.08-0.1 Hz range for very low noise (*σ* = 0-1 × 10^-6^; Fig. 8B) and for the entire noise range (Fig. 8C).

**Figure 8:**
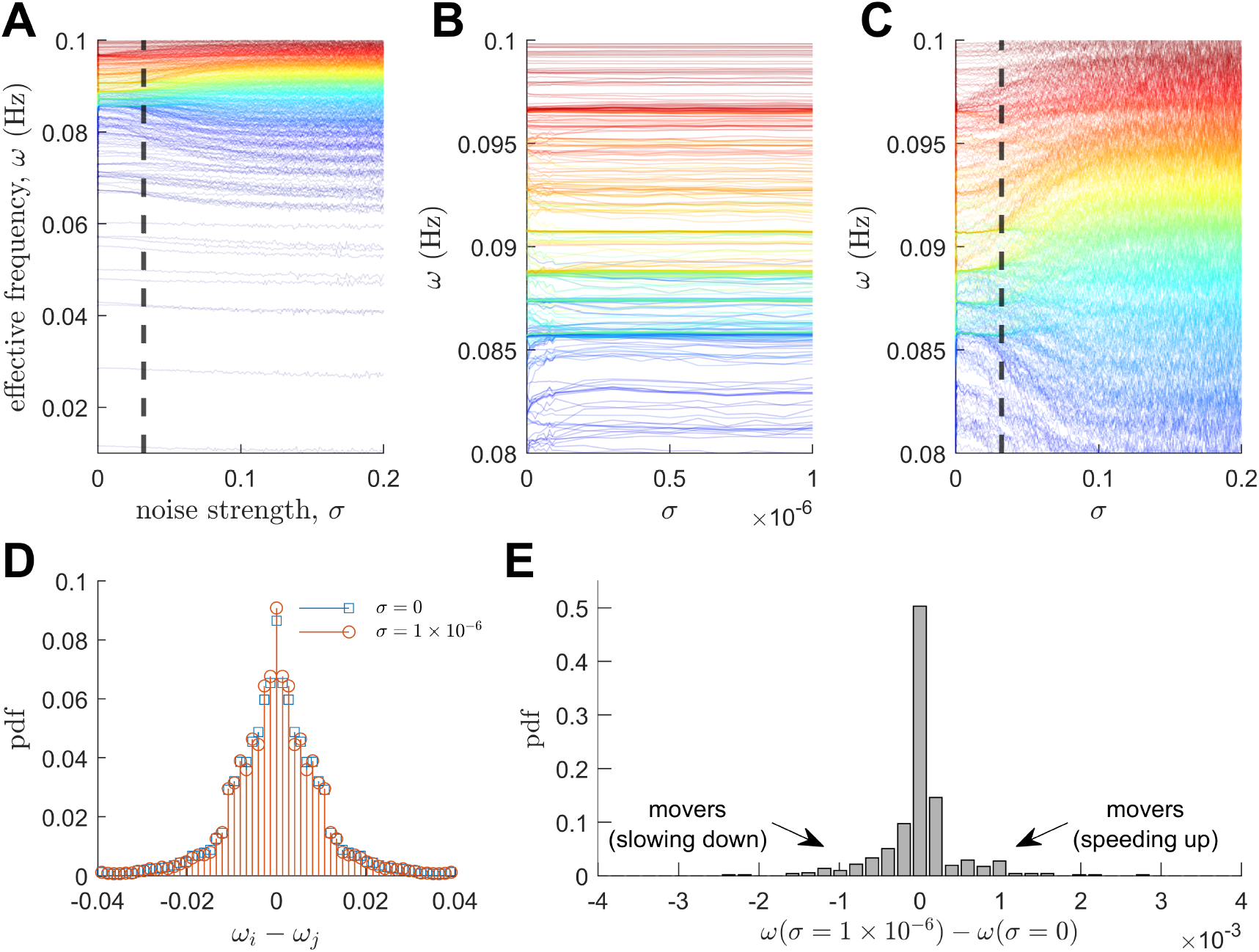
Frequency clustering. **(A)** Effective node frequency *ω* vs noise strength *σ*. **(B)** Same as panel A but for very low noise strengths (*σ* = 0–1 × 10^-6^). The y-axis is zoomed to show the range *ω* = 0.08–0.1 Hz. **(C)** Same as panel B but for the entire noise range (*σ* = 0–0.2). For panels A, B, and C, the lines are colored according to the order of natural frequencies (blue: low *ω*^0^; red: high *ω*^0^). The lines also represent ensemble averages of 50 noise realizations. For panels A and C, the dashed line represents the *σ* where Δ*S* in Fig. 4A becomes negative (i.e., desynchronization). **(D)** Pdf of internode frequency difference (*ω_i_ – ω_j_*). The blue square markers are for *σ* = 0 and the red circle markers are for *σ* = 1 × 10^-6^. **(E)** Pdf of the difference of *ω* at *σ* = 1 × 10^-6^ and *σ* = 0. This identifies the *movers,* which are nodes that speed up and slow down.

Recall from Fig. 2D that at *σ* = 0, we found that interactions between nodes alone result in the formation of several frequency clusters (i.e., alignment of frequencies), arising due to the network being posed in a state of high metastability. We see from Fig. 8B that turning on the noise, albeit at very low strengths, reorganizes the frequency clusters. That is, noise perturbs the nodes asymmetrically such that some stay at their noise-free effective frequencies and some move faster or slower to aggregate with existing clusters (e.g., the predominantly upward shift in the blue clusters in Fig. 8B), resulting in the formation of larger clusters (see higher occurrence of internode frequency differences near 0 compared to the noise-free case in Fig. 8D). These larger clusters persist from very low to moderate noise strengths (Fig. 8C), which is why synchronization increases for moderate noise (the stochastic synchronization effect). Increasing noise further, especially beyond the cut-off where Δ*S* in Fig. 4 becomes negative (represented by the dashed line in Fig. 8C), dissolves the clusters and the nodes continuously change their frequencies until they return to their natural frequencies (which agrees with the findings in Fig. S11).

Nodes that changed their frequencies in Fig. 8B to join existing frequency clusters, which we call as the *movers*, can be identified by taking the pdf of the difference of *ω* at a chosen low to moderate *σ* value (in our case, we choose *σ* =1 × 10^-6^) and *ω* at *σ* = 0, as shown in Fig. 8E. These movers play a crucial role in increasing the network synchronization (due to them making frequency clusters bigger and shifting the clusters toward the mean frequency). Interestingly, we find that several of the biggest *movers* belong the the hub network (the first 20%) defined in Fig. 4B; hence, this could be why the effect is most pronounced in that network. Moreover, we note that the frequency-clustering mechanism described above persists only when there are already several frustrated clusters to begin with (at σ = 0). Most importantly, we find that the network topologies investigated in Fig. 6C are not conducive to the creation of these frustrated clusters (Fig. S12); hence, why they cannot produce the stochastic synchronization effect.

Finally, to investigate the phenomenon based on large-scale network dynamics, we partition the connectome into 12 empirically derived functional network subdivisions of the brain [63] (Fig. S13). Then, we calculate within and between subnetwork functional connectivity *FC* by averaging the cosine correlations between all pairs of regions within the subnetworks (see Materials and Methods for details); a positive FC means that the concerned subnetworks have correlated dynamics, hinting a positive functional relation. Figures 9A–9C show that the overall qualitative pattern of *FC* is similar for the noise-free and moderate-noise cases, but abolished for the high-noise case. However, quantitatively, in general, the moderate noise strength increases functional connectivity. The increases are more apparent by measuring the change in functional connectivity Δ*FC* with respect to the baseline (i.e., the noise-free case), as shown in Fig. 9D. However, we emphasize that the changes are nonuniform across all pairs of subnetworks, with some pairs exhibiting a decrease (e.g., *FC* of MEM-AUD and SUB-MEM) or no change (e.g., *FC* of VIS-VIS and FP-SH) in functional connectivity, as shown in Fig. 9E. Moreover, a high noise strength destroys the functional connectivity of all subnetworks and the global network, ultimately leading to almost zero FC, which corroborates our previous point that the action of high noise pushes the oscillations into an incoherent random state, hence becoming uncorrelated. Moreover, in the context of these functional networks, we find that majority of the significant frequency movers in Fig. 8C (i.e., nodes that sit at the tails of the pdf) belong to DM and VIS networks.

**Figure 9:**
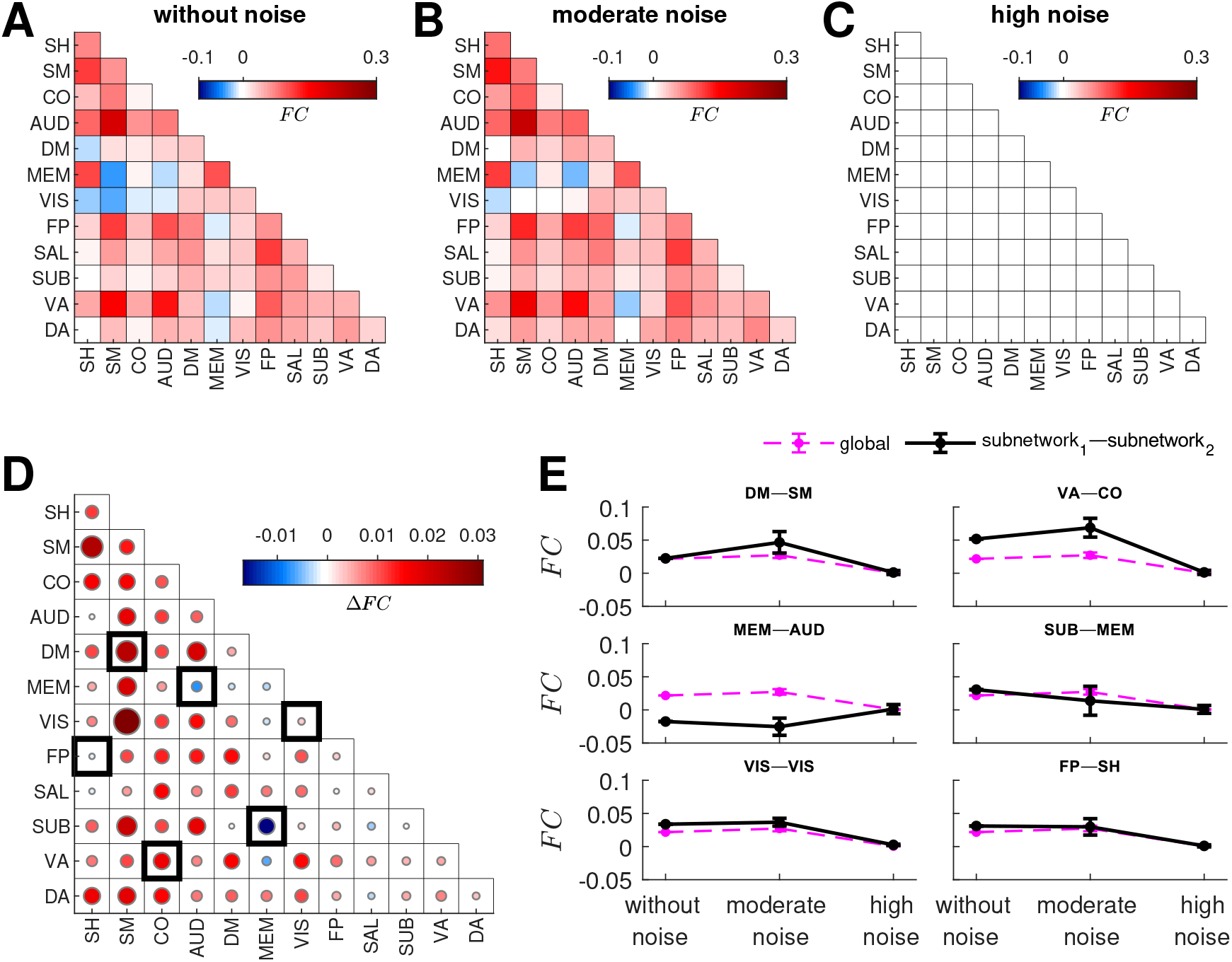
Functional connectivity of subnetworks for various noise strengths. **(A)** Functional connectivity *FC* within and between subnetworks for dynamics without noise for one realization. **(B)** Same as panel A but with moderate noise. **(C)** Same as panel A but with high noise. **(D)** Change in functional connectivity Δ*FC* from dynamics without noise to dynamics with moderate noise. Exemplar *FC* of subnetworks enclosed by the solid boxes are shown in panel E. **(E)** Exemplar *FC* between subnetworks (black solid lines and labeled). The magenta dashed lines represent the global *FC* of the whole connectome. The markers represent ensemble averages of 50 noise realizations and the vertical lines represent standard deviations.

Overall, these results show that stochastic synchronization manifests on the human connectome as a reconfiguration of functional integration between regions and subnetworks: noise can merge functional modules through increased phase and frequency clustering and alter the functional connectivity of the connectome’s subnetworks.

### 2.6. Synthetic models

In Sec. 2.5, we showed that one of the necessary ingredients for stochastic synchronization to occur is for the noise-free network to be in a state where noise allows the system to access a less-frustrated and more-synchronized state. Hence, we next investigate a scenario with strongly frustrated frequencies such that noise can effectively speed up slow oscillations and slow down fast oscillations, leading to increased synchronization.

We construct an instructive synthetic model with a fully connected (all-to-all) network and frustrated distinct natural frequencies (i.e., linearly spaced from 0.01 to 0.1 Hz; the same frequency range used above). Moreover, the dynamics are placed at a highly metastable state at zero noise (similar to the method in Fig. 2). We find that this network can strongly produce the effect (Fig. S14A) via a frequency-clustering mechanism. Low to moderate noise noise enhances synchronization as the slowest nodes move faster and the fastest nodes move slower towards a mean field (Fig. S14B). In the limit of strong noise, the nodes tend to follow their natural frequencies (similar to the results in Fig. 8). These results demonstrate that stochastic synchronization can indeed occur in simple networks provided they intrinsically have frustrated frequency distributions.

The effect also occurs for two connected oscillators (Figs S15A and S15C; with natural frequencies set to 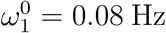 and 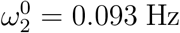 to mimic nodes on either side of the mean-ensemble frequency of the human connectome model), even though it is well-known that noise monotonically degrades frequency locking (Fig. S15B) [42]. In this two-oscillator case, the effect is produced via a phase-clustering mechanism, where noise assists in producing more phase slips that increase synchronization (see [61] for a thorough analysis).

Finally, we study another synthetic model with a random modular network (Fig. S16A; network with four modules) and frustrated natural frequencies (the natural frequencies of each module are set to *ω*^0^ = 0.01, 0.04, 0.07, and 0.1 Hz). By construction, this synthetic model induces the frequency clustering mechanism discussed in Fig. 8 and can also produce the effect as predicted (Fig. S16B).

Overall, all our results and analyses demonstrate that stochastic synchronization is partly driven by a frequency-clustering mechanism and partly by a phase-clustering mechanism, which are both embodied in our human connectome network with an anatomically-based hierarchy of natural frequencies because of the existence of dynamically frustrated states.

## 3. Discussion

In this work, we used a computational brain network model to investigate the synchronization of cortical oscillations on a large-scale human connectome. We revealed the existence of a counterintuitive phenomenon (i.e., stochastic synchronization) where the addition of disorder (random noise) to the brain can yield a more ordered (synchronized) state. We then teased apart and explained the network and dynamical origins of this phenomenon.

One of the advantages of the Kuramoto-based brain network model is its low complexity, relying on only two parameters; i.e., the coupling strength *c* and noise strength σ. This is why the Kuramoto literature has a wide range of available analytical results for well-defined networks (e.g., random networks) and frequency distributions (e.g., Gaussian or Lorentzian). However, since these analytical results cannot be straightforwardly applied to more biological networks and frequency distributions (such as ours) in the presence of noise [64], we extensively investigated via numerical methods how network dynamics change with respect to the two parameters *c* and *σ*. Varying the coupling strength parameter alone, which scales the overall influence of activity between connected regions, can produce transitions between incoherent dynamics and coherent global synchronization patterns [8, 12]. At a critical coupling strength, network coherence is highly metastable such that the network spontaneously and transiently visits incoherent and coherent states. Our investigations focused on this near-critical regime because it has consistently been found to be where network models best fit neuroimaging data [29, 30, 31, 32, 57, 33]. As a side note, shifts in the location of the critical point or being tuned to coupling strengths farther away from the critical point may be a fingerprint of suboptimal brain function or neurological and psychiatric disorders (e.g., [65, 66]), which could be an interesting avenue to investigate in the future. Our results reveal a new role for near-critical metastable dynamics in stochastic synchronization. Moreoever, we extend earlier results on synchronization transitions in Kuramoto models on the connectome to the case of neurobiologically-realistic heterogeneity in local oscillator frequencies.

It is usually expected that the behavior or function of a physical system should deteriorate in the presence of stochasticity such as noisy inputs. However, we showed that this is not the case for the human connectome. We demonstrated that stochastic synchronization emerges, where moderate levels of noise can synchronize global and local dynamics on the network, but only around the critical coupling strength (Fig. S3). Moreover, we found that structural hubs can benefit the most from the addition of noise, exhibiting much higher increases in synchronization as compared to nonhubs. Hubs are well known to be critically important for brain network integration, enabling efficient neural signaling and communication to foster complex cognitive functions [67, 23, 68, 55]. Our results imply that hubs are the regions most able to facilitate stochastic synchronization. This suggests a new role for hubs in orchestrating brain network integration in the face of noisy inputs.

Given that the brain is constantly bombarded with stochastic influences across multiple spatiotemporal scales [69], stochastic synchronization offers an explanation as to how the brain, as a whole, can robustly adapt to and utilize randomness to achieve normal function and/or to engender emergent behavior. By systematically evaluating the contributions of network topology and local node characteristics to the emergence of stochastic synchronization, we found strong evidence for the important role of hierarchy and heterogeneity in the brain. In particular, we uncovered that (i) the hierarchy of timescales imposed by intrinsic neuroanatomy [25, 56] and (ii) the heterogeneous complex topology of the connectome with hubs and nonhubs positioned across the brain [21, 22] provide a substrate for the brain to attune to stochastic influences and maintain/enrich its overall dynamics. Moreover, we emphasize that the effect emerges due the complex synergy of the brain’s network topology and hierarchy of natural frequencies via the creation of dynamically frustrated states. Previous studies have shown that atypical neural timescales [70], fragility and volatility of hubs in the connectome [55], and interactions between noise and hubs [71] are associated with neuropsychiatric disorders. Hence, these dynamical and topological perturbations in brain disorders and diseases likely also manifest in the extent of the brain network’s stochastic synchronization, which could be measured using the framework developed in this study. In fact, it would be interesting to investigate in the future whether the optimal working point (corresponding to values of *c* and *σ* that better fit fMRI data) changes for each individual, over time (e.g., dynamic functional connectivity [62, 72]) different brain states (e.g., resting vs task, sleep vs wake), or clinical conditions (e.g., healthy vs diseased, young vs old). It is also plausible that the optimal set point varies dynamically in line with internal (e.g., neuromodulation [73]) and environmental factors, which could ultimately influence the brain’s ability to support diverse functional processing [74].

We then revealed that the action of a moderate level of noise, where stochastic synchronization thrives, is three-fold. First, moderate noise entrains node phases to align with each other, resulting in fewer phase clusters with predominantly bigger sizes. That is, increased integration within functional modules (clusters of regions). This translates to changes in the level of synchronization, with a stronger stochastic synchronization effect arising from a minimal number of clusters. Second, moderate noise pushes some nodes to move faster or slower to temporally lock their oscillations with other nodes. Thereby resulting in large frustrated frequency clusters that eventuate higher levels of synchronization. Third, moderate noise alters the functional connectivity between subnetworks in the brain, either boosting or diminishing it. These actions of noise could in principle be tested experimentally. For example, pharmacological manipulations that increase/decrease neuronal noise could be used to investigate changes in integration and functional network architecture. These manipulations could include the administration of psychedelic drugs, such as LSD and psilocybin, that target underlying neuroreceptors and have been hypothesized to increase the entropy of the brain, resulting in a more disordered state of consciousness [75]. In fact, recent studies have hinted the ability of these psychedelics to influence reorganization of functional brain networks [76, 77] but the actual mechanisms of this action are unclear. Our work may provide insights into these mechanisms. However, the precise protocols to test the effects of these drugs in the future must be carefully considered because they could potentially simultaneously affect multiple physiological properties (e.g., neuronal excitability, coupling, frequencies of oscillation, and noise within local circuits). Our study also has potential clinical applications. The stimulant methylphenidate has been linked to decreased neuronal noise in the treatment of attention-deficit/hyperactivity disorder [78]. Moreover, our measurements of network response (i.e., coherence, synchronization, and metastability) can be easily adopted by future clinical studies, for example, in diagnostic or treatment monitoring purposes such as tracking of network excitability in response to antiepileptic drugs [79].

Finally, we emphasize that the formulation of this study is general because at the core of our computational brain network model is the paradigmatic Kuramoto-type phase oscillator, which can conveniently represent coupled oscillatory units in many neural and nonneural systems. Hence, the stochastic synchronization phenomenon demonstrated in this study may be generalized to other systems (including outside neuroscience) as long as the system’s network topology and intrinsic dynamics exhibit some sort of heterogeneity and hierarchy, as discussed above. Moreover, the effect could also be explored using more complex and realistic neural mass models [30, 80, 34].

In summary, our study demonstrates the use of a simple mesoscopic brain network model in providing a mechanistic understanding of the synchronization dynamics of cortical oscillations. It allowed us to easily generate diverse patterns of synchronization on the human connectome that may represent various functional states of the brain. Moreover, using this approach, we revealed that when the connectome operates near a critical coupling, the novel phenomenon of stochastic synchronization emerges driven by the topological and dynamical properties of the brain. This can be exploited to study its relation to normal and/or pathological brain function, which would be an exciting avenue to test in the future.

## Materials and Methods

### Connectomic data

The whole-brain human structural connectome was derived from 75 healthy participants (aged 17-30 years, 47 females) using diffusion weighted MRI [52, 34]. MRI data were acquired using a Philips 3 T Achieva Quasar Dual MRI scanner with a single-shot echo-planar imaging (EPI) sequence (TR = 7767 ms, TE = 68 ms). High-resolution whole-brain fiber tracks were generated via a probabilistic streamline algorithm with constrained spherical deconvolution, as implemented in the MRtrix software [81]. Undirected structural connectivity matrices for all participants were constructed from densely seeded tractography and parcellated into *N* = 513 cortical and subcortical regions of approximately uniform size [82]. The weights of the matrices represent the number of streamlines linking each pair of regions divided by the streamline length. Finally, to filter out idiosyncratic variations, the final connectivity matrix in Fig. 1 refers to a group-averaged connectome. Further details of MRI scanner properties, data preprocessing, parameters of the tractography algorithm, and parcellation are provided in our previous study [52].

For replication of results, minimally preprocessed diffusion weighted MRI data from 2 unrelated healthy young adult participants (participant IDs: 100206, 100307) were obtained from the Human Connectome Project (HCP) [53, 83]. The data were acquired on a customized Siemens Magnetom Skyra 3T MRI system according to the following parameters: TR = 5520 ms, TE = 89.5 ms, 3 diffusion-weighted shells (b = 1000, 2000, and 3000 s/mm^2^), 145×145 matrix, 174 slices, and 1.25× 1.25× 1.25 mm^3^ voxel size. The diffusion images were further processed using the MRtrix software [81] by applying bias-field correction and multishell multi-tissue constrained spherical deconvolution to model white matter, gray matter, and cerebrospinal fluid. For each HCP participant, tractograms were generated using 10 million probabilistic streamlines, 2nd-order Integration over Fiber Orientation Distributions algorithm (iFOD2), anatomically-constrained tractography (ACT) [84], dynamic seeding [85], backtracking, streamline lengths of 5-250 mm, and spherical-deconvolution informed filtering of tractograms (SIFT2) [85]. Each participant’s tractogram was used to create connectomes parcellated into two commonly used FreeSurfer atlases: Desikan-Killiany Atlas (*N* = 164 cortical and subcortical regions) [86] and Destrieux Atlas (*N* = 84 cortical and subcortical regions) [87]. The weights of the connection matrices represent the streamline density, with each streamline multiplied by the appropriate weighting factor generated by the SIFT2 method, scaled by the inverse of the streamline length.

### Simulation details

The simulations were performed in MATLAB (version 2018b, MathWorks Inc.) by numerically integrating the stochastic differential equation in Eq. (1) via the Euler-Maruyama scheme with a timestep of *dt* = 0.25 *s* for a total time of 15000 s. We verified that using a different integration scheme (Heun) and shorter timesteps (i.e., *dt* = 0.05, 0.1 s) does not change the results of the study (Fig. S17). Unless otherwise stated, the results refer to ensemble averages of either 30 different initializations (i.e., initial conditions) or 50 noise realizations using the same initial condition.

### Measures of network dynamics

The phase solutions of Eq. (1) were wrapped into the interval [0, 2*π*], where 0 maps to 0, positive multiples of 2*π* map to 2*π*, and negative multiples of 2*π* map to 0. To quantify instantaneous collective dynamics at time *t*, we used the order parameter 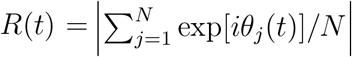, which measures global network coherence and varies between 0 (fully incoherent state) and 1 (fully coherent state). To ensure that our results only captured steady-state dynamics, we discarded transients and only kept data for the time period *T* = 5000-15000 s. All subsequent measurements in this study were performed on this steadystate time period.

Network synchronization *S* was calculated by taking the time average of *R*(*t*), while network metastability *M* was calculated by taking the standard deviation of *R*(*t*). To highlight changes in the synchronization of a system with noise strength *σ*, the percent change in network synchronization Δ*S* was calculated as [*S*(*σ*) — *S*(*σ* = 0)] /*S*(*σ* = 0) × 100. To quantify changes in the nodes’ oscillation frequencies, we calculated the effective frequency of node *i* as 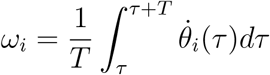, where *T* is the steady-state time period defined above [58]. We also measured the correlation of oscillations between nodes *i* and *j* called functional connectivity *FC* calculated as *FC* = 〉cos(*θ_i_* — *θ_j_*)), where the brackets denote steady-state time average; *FC* varies from —1 (anticorrelated), 0 (uncorrelated), and 1 (correlated). The metric was extended to capture the functional connectivity between subnetworks of the connectome by taking the average FC among all pairs of nodes in the subnetworks.

### Frequency distributions

The other distributions of natural frequencies in Fig. 5 were: (i) Dirac-delta (homogeneous); (ii) random uniform (rand-uniform); (iii) random Gaussian (rand-gaussian); and (iv) random Lorentzian (rand-lorentzian). The distributions were bounded to the frequency bandwidth of fMRI (i.e., 0.01-0.1 Hz) similarly to Eq. (2). Distribution (i) comprised frequencies that were all set to *ω*_mean_ = 0.055 Hz. Distribution (ii) comprised uniform-distributed random frequencies. Distribution (iii) comprised Gaussian-distributed random frequencies centered at *ω*_mean_ with a standard deviation of *ω*_mean_/5. Distribution (iv) comprised Lorentzian-distributed random frequencies with a median of *ω*_mean_ and a half width at half maximum of *ω*_mean_/5.

### Network topologies

The other network topologies used in Fig. 6 were: (i) fully connected; (ii) weight-preserving random network, randomizing 50% of the connections; (iii) weight-preserving random network, randomizing 100% of the weights; (iv) geometry-preserving random network, preserving the node strengths; and (v) geometry-preserving random network, preserving the node-strength sequence. Topology (i) was a fully-connected network with all-to-all connections and weights equal to the average weight of the human connectome. Topology (ii) was a surrogate of the human connectome with 50% of the weights randomly shuffled. Topology (iii) was a surrogate of the human connectome with 100% of the weights randomly shuffled. Topology (iv) was a surrogate of the human connectome with randomly shuffled weights but preserving the weight-distance relationship and the distribution of node strengths, which means that the surrogate network has hubs with the same node strengths as those in the empirical connectome but now positioned randomly. Topology (v) was a surrogate of the human connectome with randomly shuffled connections but preserving the weight-distance relationship and the node strengths in their original order, which means that all nodes have the same strengths and positions as those in the empirical connectome. The algorithms to create topologies (ii)-(v) were taken from [52, 55]. To quantitatively compare the structure of these topologies, we calculated their small-world propensity [88], which is a recently developed graph-theory measure that quantifies the extent to which a weighted network displays small-world characteristics inspired by [89].

### Phase-clustering algorithm

Phase clustering was analyzed using a density-based clustering algorithm DBSCAN [90]. The advantage of DBSCAN against other popular community detection algorithms, such as the Louvain and k-means algorithms, is that it is robust to outliers and does not need an *a priori* value for the number of clusters. The algorithm requires two parameters to be defined: cluster radius and minimum cluster size. The cluster radius *ϵ* specifies the maximum distance between nodes for them to be considered as part of a cluster. Since the nodes were placed on a unit circle (see Fig. 7A), the cosine distance was used, which is one minus the cosine of the phase difference between nodes. The minimum cluster size minPts specifies the minimum number of nodes needed to form a cluster. We set *ϵ* = 0.0011 and minPts = 10 to optimize the number of clusters formed (see optimization in Fig. S18), but we verified that changing the values of these parameters does not change the results of the study.

### Null models

Two null models, i.e., uncoupled and random, were generated for comparing the synchronization and phase-clustering statistics resulting from the high-noise case in Fig. 7. The uncoupled model was generated using our brain network model but with coupling strength set to *c* = 0, which effectively simulated a network with disconnected nodes and node phases evolving independently. On the other hand, the random model was generated by randomly choosing node phases from a uniform distribution [0, 2π], disregarding any kind of structured dynamics.

### Statistical analysis

Pair-wise differences in the synchronization and phase-clustering statistics in Fig. 7 were tested using a one-way ANOVA and corrected for multiple comparisons using the Bonferroni-Holm procedure.

### Code availability

MATLAB codes to perform sample simulations and generate the figures of this study are available at https://github.com/brain-modelling-group/stochastic-sync.

## Acknowledgments

This work was supported by the Australian National Health and Medical Research Council (Project Grants 1145168 and 1144936, and Fellowship 1110975). HCP data used for replication were provided by the Human Connectome Project, WU-Minn Consortium (Principal Investigators: David Van Essen and Kamil Ugurbil; 1U54MH091657) funded by the 16 NIH Institutes and Centers that support the NIH Blueprint for Neuroscience Research, and by the McDonnell Center for Systems Neuroscience at Washington University.

## Supplementary Material

**Figure S1:**
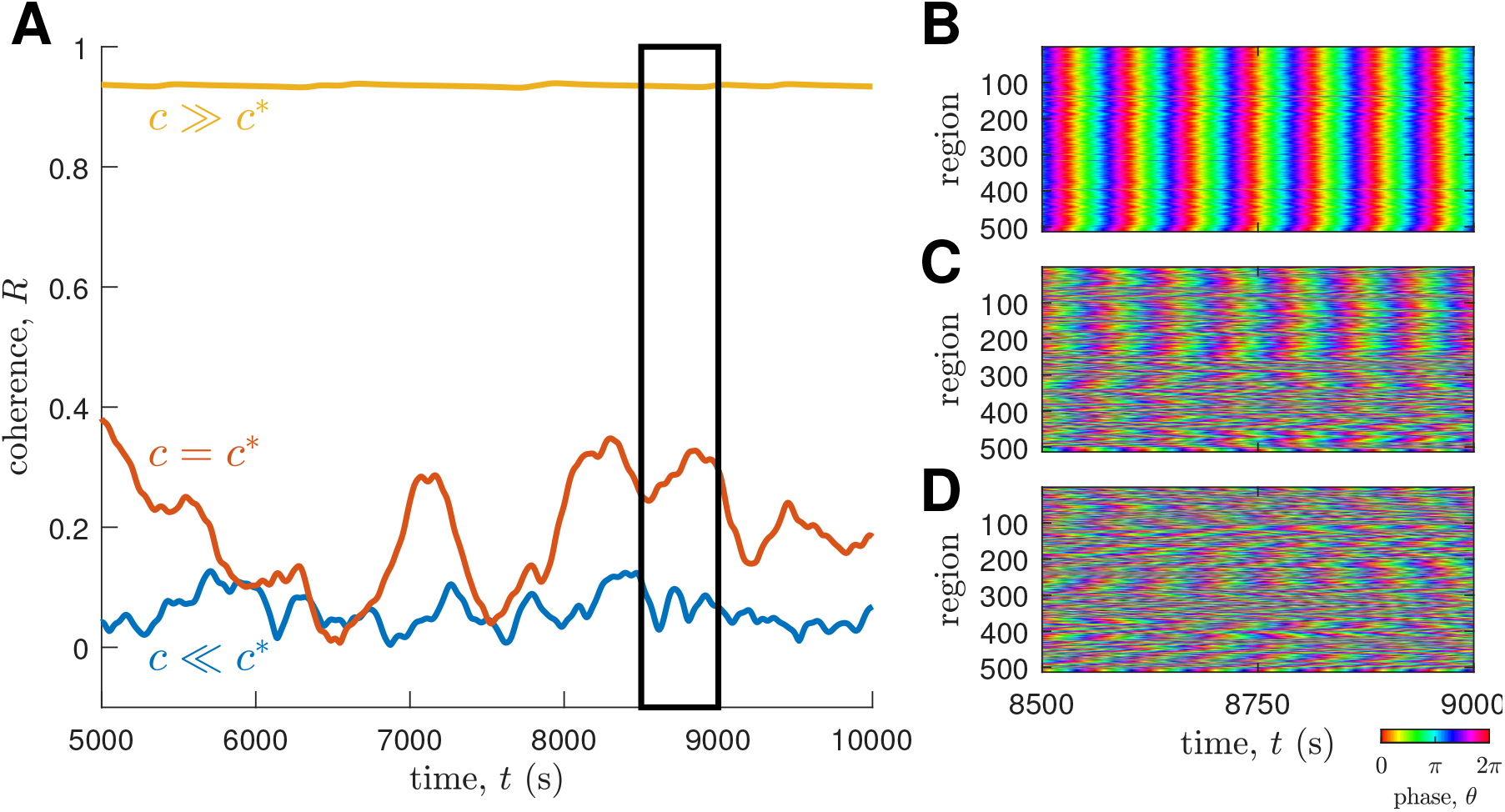
Network coherence and spatiotemporal dynamics for various coupling strengths. **(A)** Time evolution of network coherence *R* for *c* ≫ *c*\* (yellow), *c* = c* (red), and *c* ≪ c* (blue). **(B)** Local phase dynamics for *c ≫ c*\*. **(C)** Same as panel B but for *c = c*\*. **(D)** Same as panel B but for *c ≪ c*\*. For panels B, C, and D, the dynamics shown are within the time window highlighted by the black solid box in panel A.

**Figure S2:**
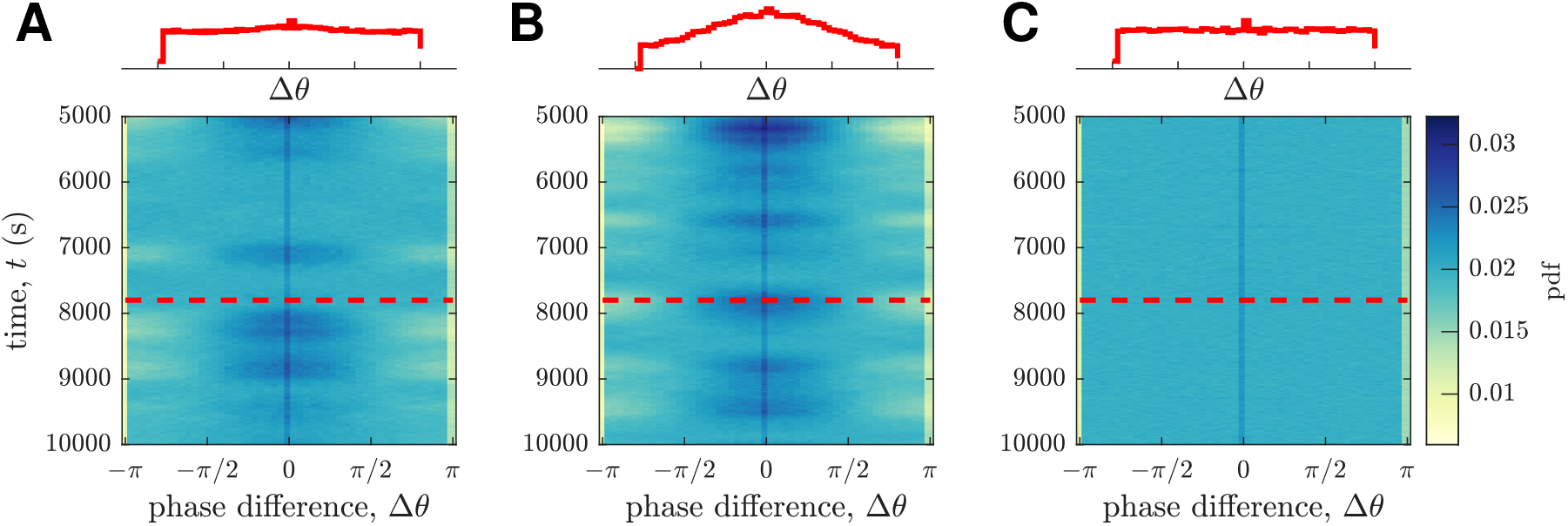
Time evolution of the distribution of phase differences for various noise strengths. **(A)** Pdf of phase difference Δ*θ* for dynamics without noise. **(B)** Same as panel A but with moderate noise. **(C)** Same as panel A but with high noise. The pdf at *t* = 7800 *s* highlighted by the red dashed line is shown above the panels.

**Figure S3:**
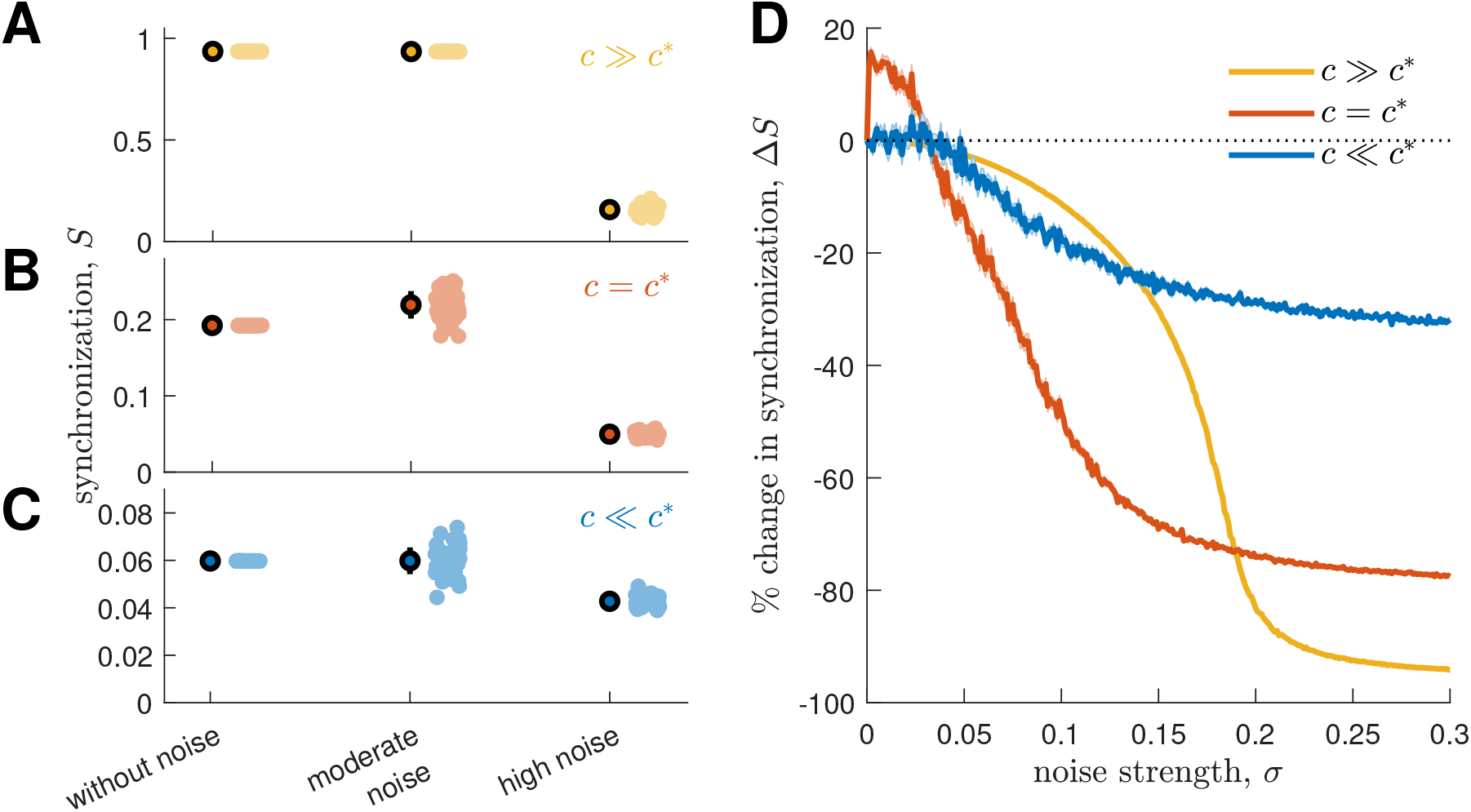
Synchronization for various coupling and noise strengths. **(A)** Synchronization *S* for *c ≫ c*\*. **(B)** *S* for *c = c*\*. **(C)** *S* for *c ≪ c*\*. For panels A, B, and C, the cloud of points represent 50 noise realizations, the thick markers represent ensemble averages of all noise realizations, and the vertical lines represent standard deviations. **(D)** Percent change of network synchronization Δ*S* vs noise strength *σ* (yellow: *c ≫ c*\*; red: *c* = *c*\*; yellow: *c ≪ c*\*). The solid lines represent ensemble averages of 50 noise realizations and the shaded areas represent the standard errors of the means.

**Figure S4:**
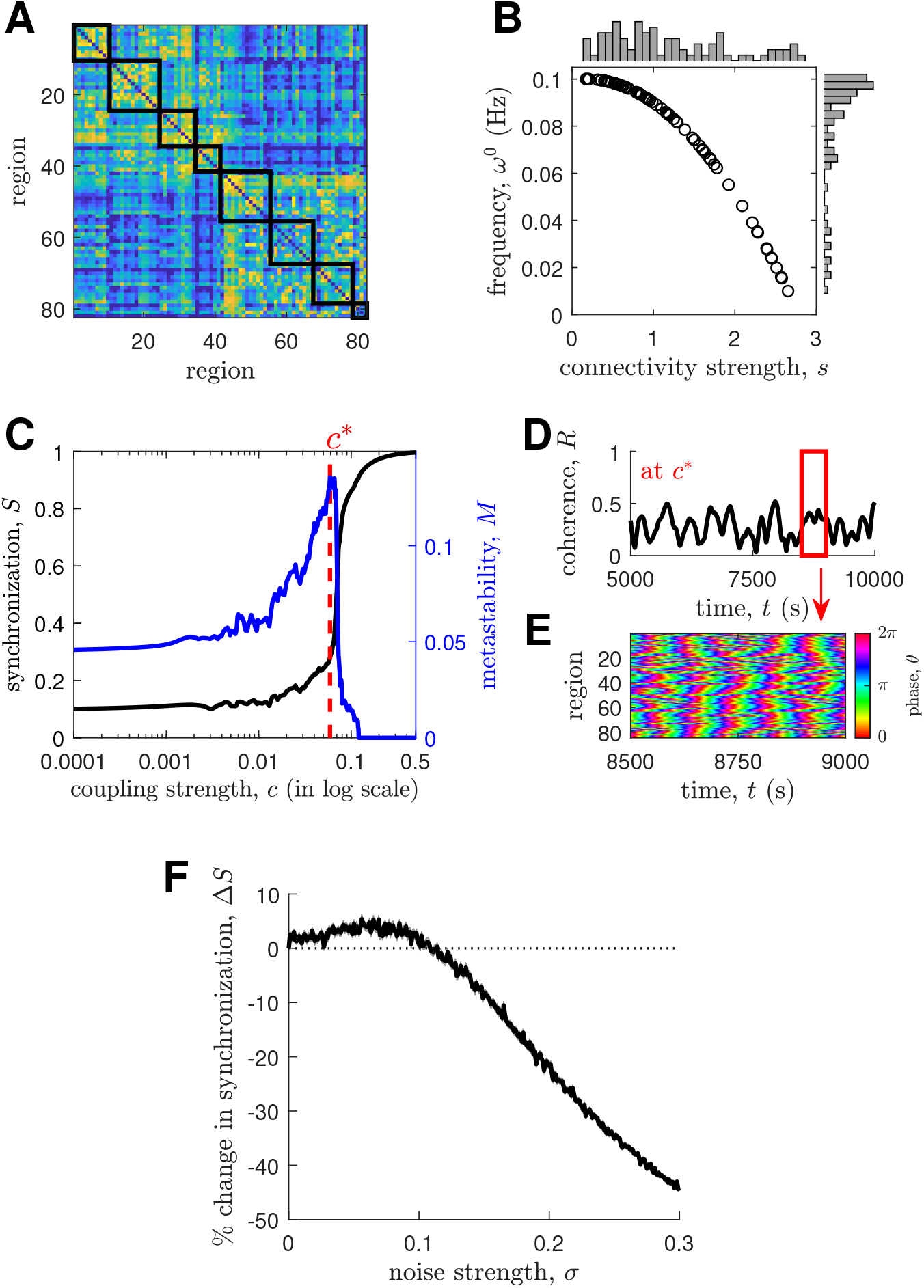
Replication of results on HCP participant 100206’s connectome with *N* = 84 regions. **(A)** Connectivity matrix with the solid boxes denoting modules. Natural frequency of oscillation *ω*^0^ as a function of connectivity strength *s*. The distributions of *s* and *ω*^0^ are shown above and to the right of the main panel, respectively. **(C)** Network synchronization *S* and metastability *M* vs coupling strength *c*. The red dashed line highlights the critical coupling strength *c*\* where *M* is maximum. The solid lines represent ensemble averages of 30 initializations. **(D)** Time evolution of network coherence *R* at *c*\*. **(E)** Local phase dynamics within the time window highlighted by the red solid box in panel D. **(F)** Percent change of network synchronization Δ*S* vs noise strength *σ* at *c*\*. The solid line represents an ensemble average of 50 noise realizations and the shaded area represents the standard error of the mean.

**Figure S5:**
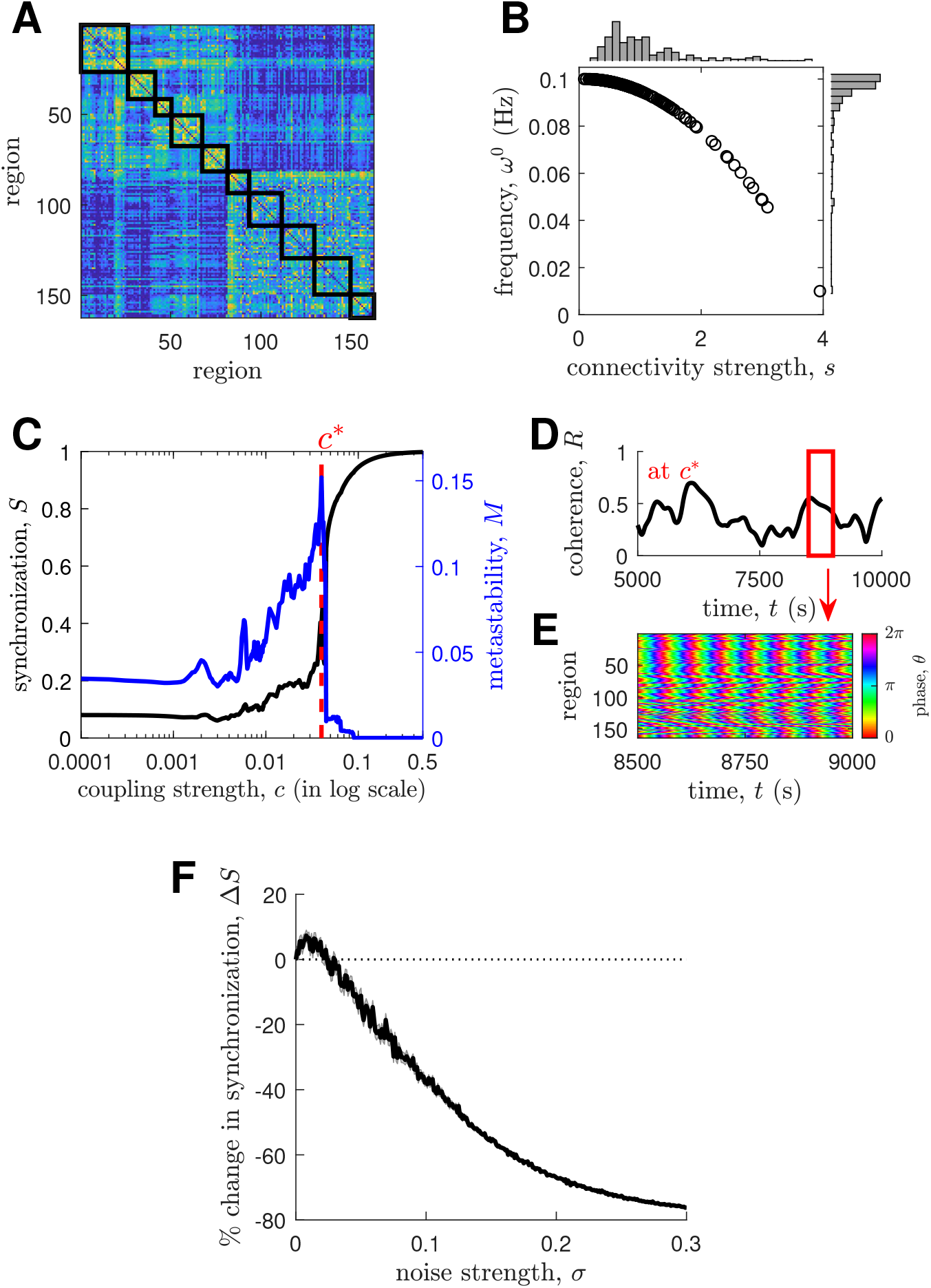
Replication of results on HCP participant 100206’s connectome with *N* = 164 regions. The details of each panel are the same as those in Fig. S4.

**Figure S6:**
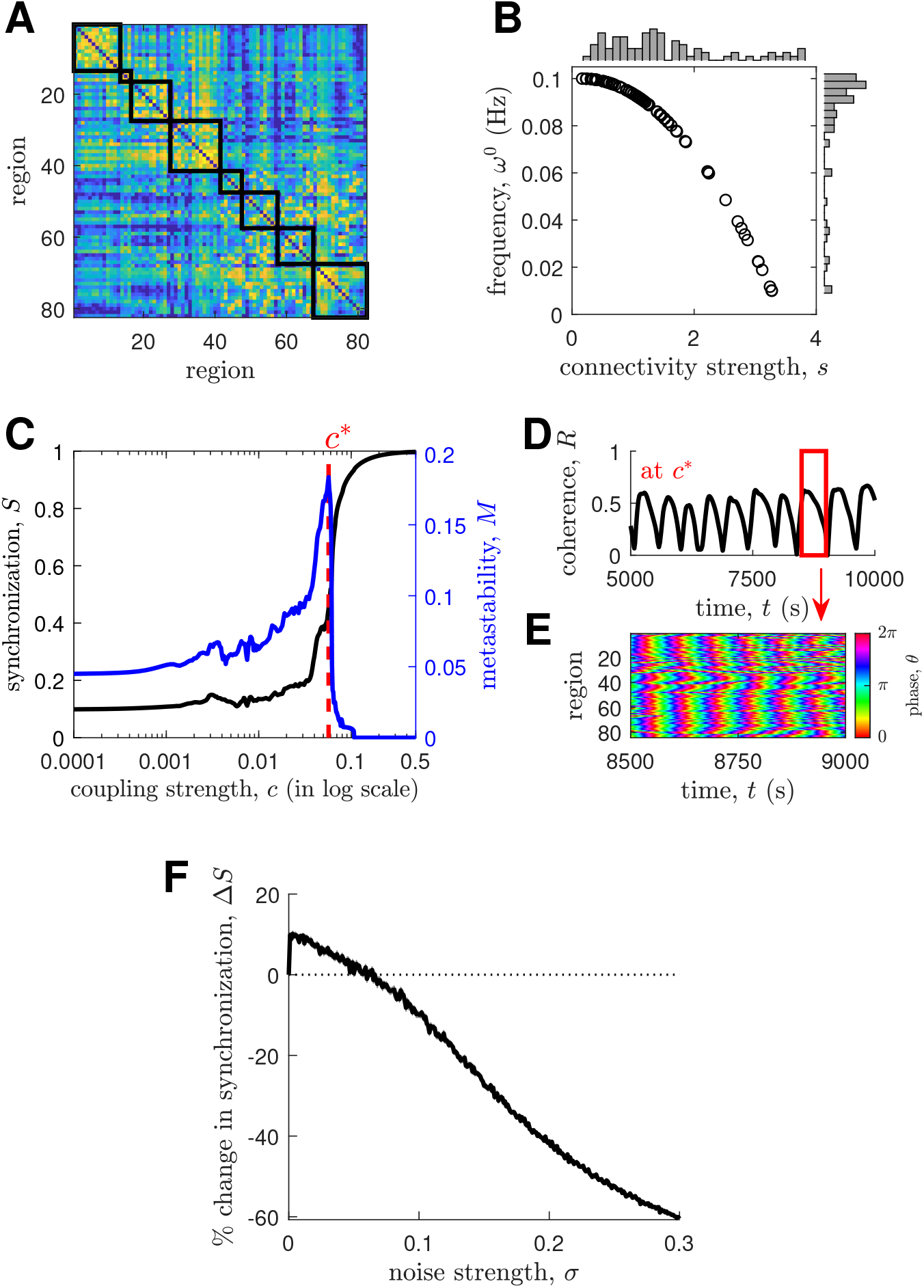
Replication of results on HCP participant 100307’s connectome with *N* = 84 regions. The details of each panel are the same as those in Fig. S4.

**Figure S7:**
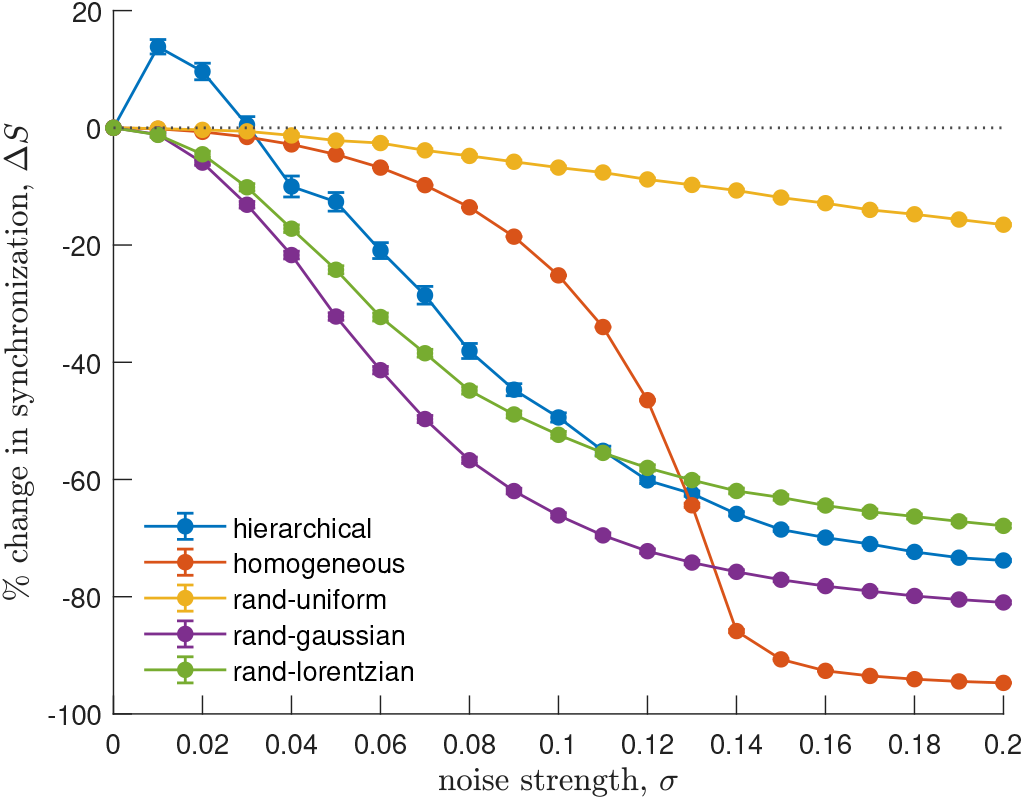
Percent change of network synchronization Δ*S* vs noise strength *σ* for different frequency distributions. The markers represent ensemble averages of 500 realizations of the frequency distributions and the vertical lines represent the standard errors of the means.

**Figure S8:**
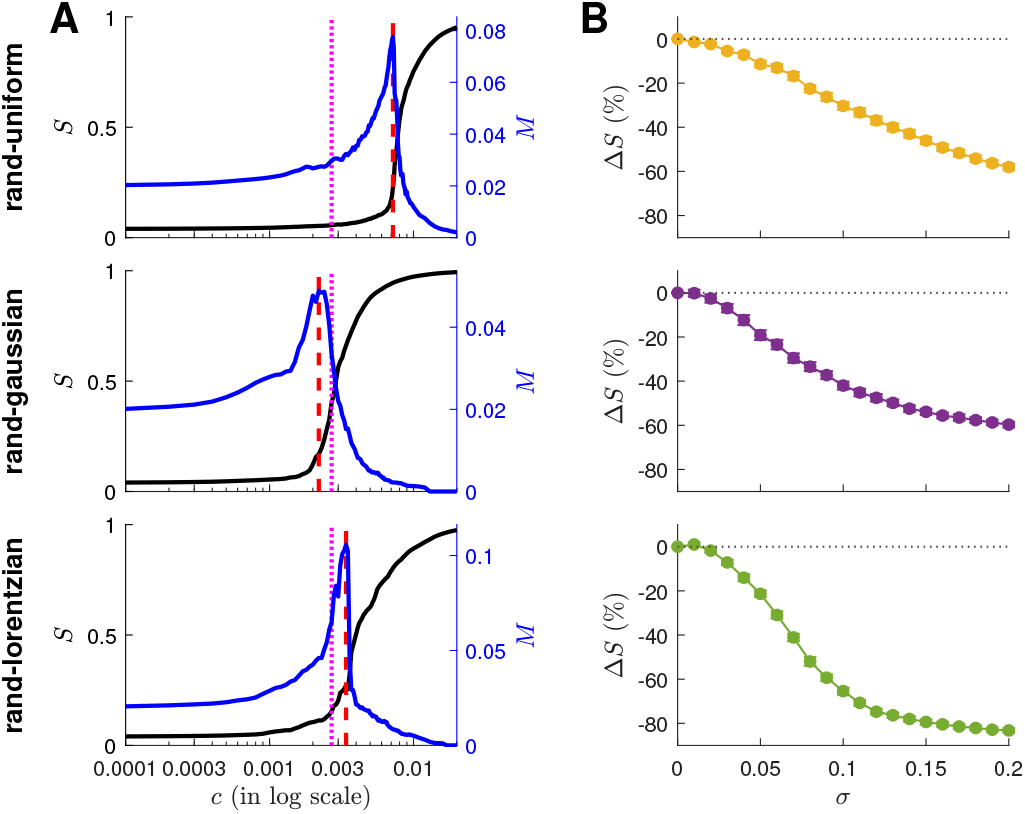
Network dynamics tuned to the critical coupling strength for different frequency distributions. **(A)** Network synchronization *S* and metastability *M* vs coupling strength *c*. The red dashed line highlights the critical coupling strength *c*\* where *M* is maximum. The magenta dotted line highlights the original critical coupling strength in Fig. 1D for ease of comparison. The solid lines represent ensemble averages of 30 initial conditions. **(B)** Percent change of network synchronization Δ*S* vs noise strength *σ*. The markers represent ensemble averages of 50 realizations of the frequency distributions and the vertical lines represent the standard errors of the means. Each row shows the results for the distribution labeled on the left.

**Figure S9:**
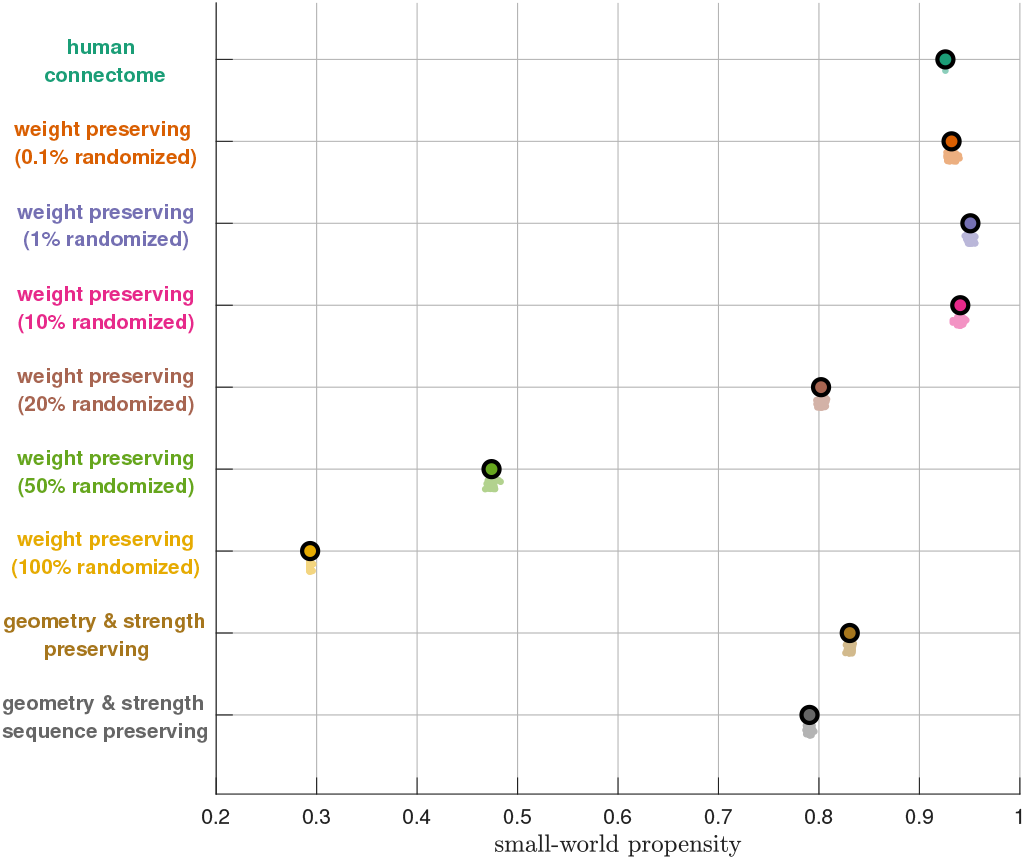
Small-world propensity of different network topologies (excluding the fully connected network). The thick markers represent ensemble averages of 50 surrogates of the network topologies and the clouds of points represent all the surrogates.

**Figure S10:**
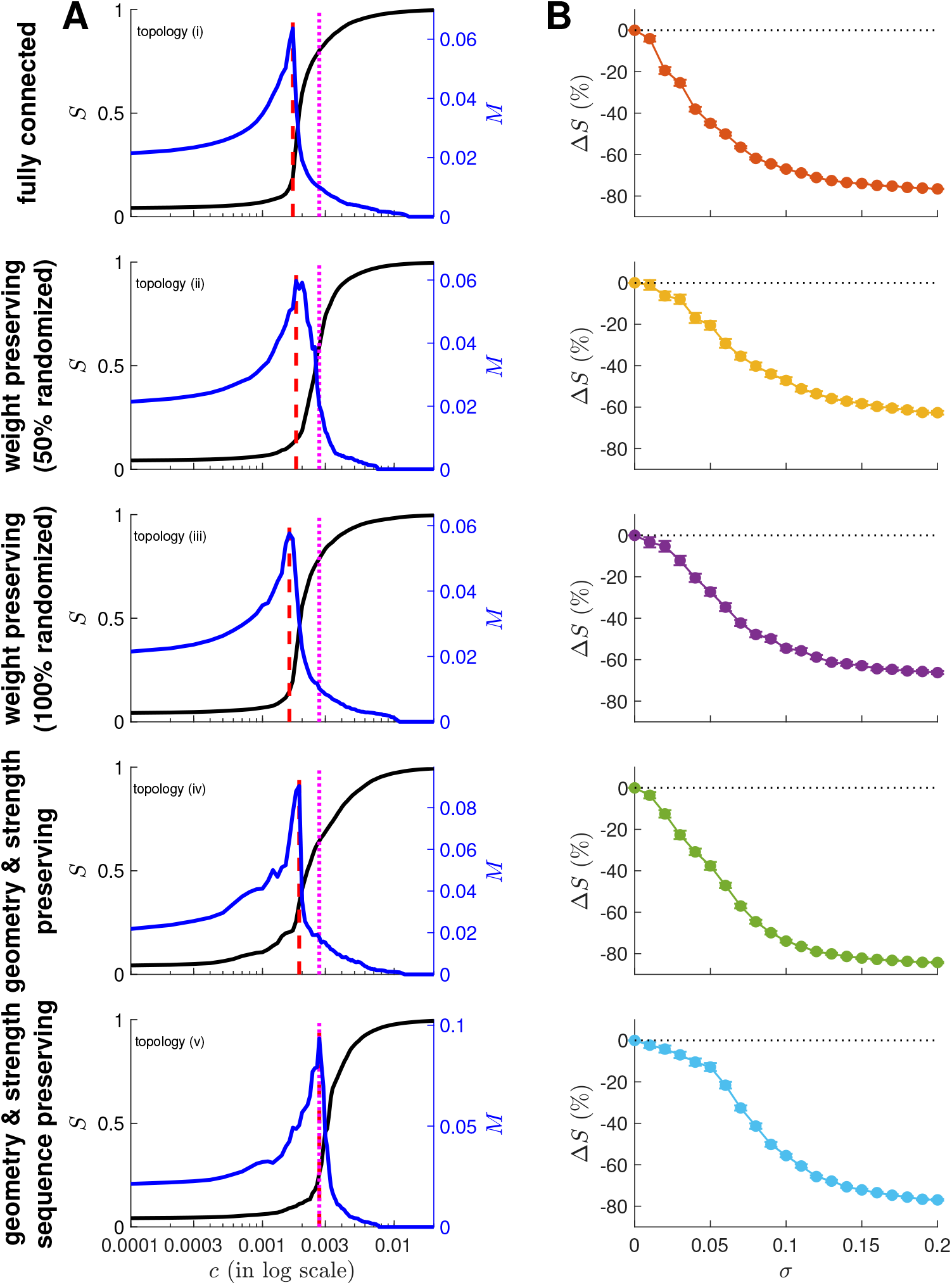
Network dynamics tuned to the critical coupling strength for different network topologies. The details of each panel are the same as those in Fig. S8. Each row shows the results for the network topology labeled on the left.

**Figure S11:**
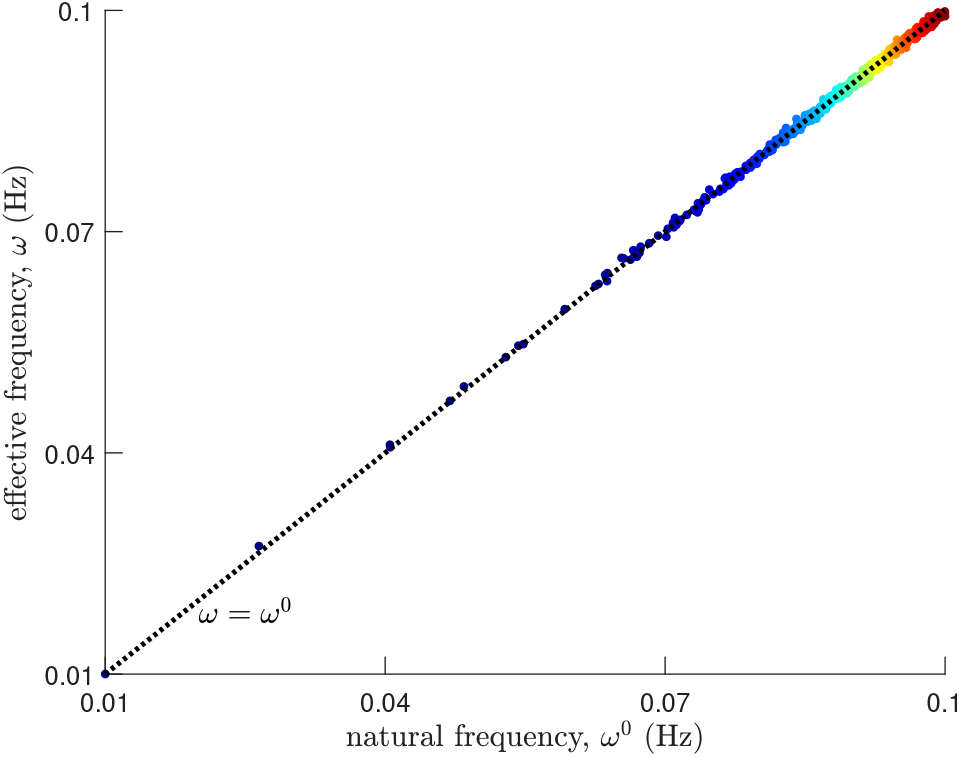
Effective node frequency *ω* as a function of natural frequency *ω*^0^ at a high noise strength. The markers are colored according to the order of natural frequencies (blue: low *ω*^0^; red: high *ω*^0^). The dashed line represents *ω* = *ω*^0^.

**Figure S12:**
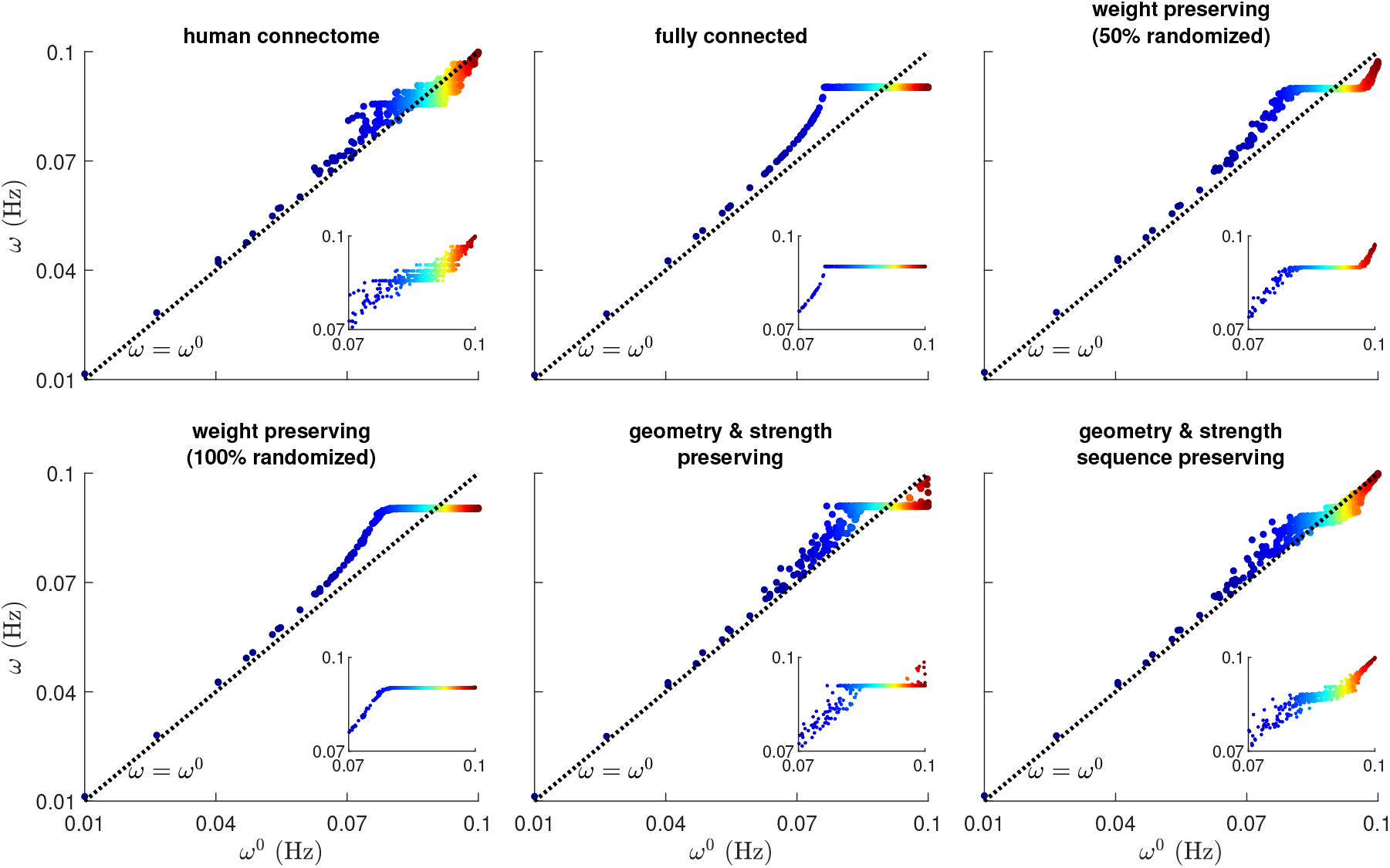
Effective node frequency *ω* as a function of natural frequency *ω*^0^ at the noise-free case for different network topologies. For all panels, the markers are colored according to the order of natural frequencies (blue: low *ω*^0^; red: high *ω*^0^). The dashed lines represent *ω* = *ω*^0^. The insets show a zoomed version of the main panels, restricting the frequencies from 0.07-0.1 Hz (for both *ω*^0^ and *ω*).

**Figure S13:**
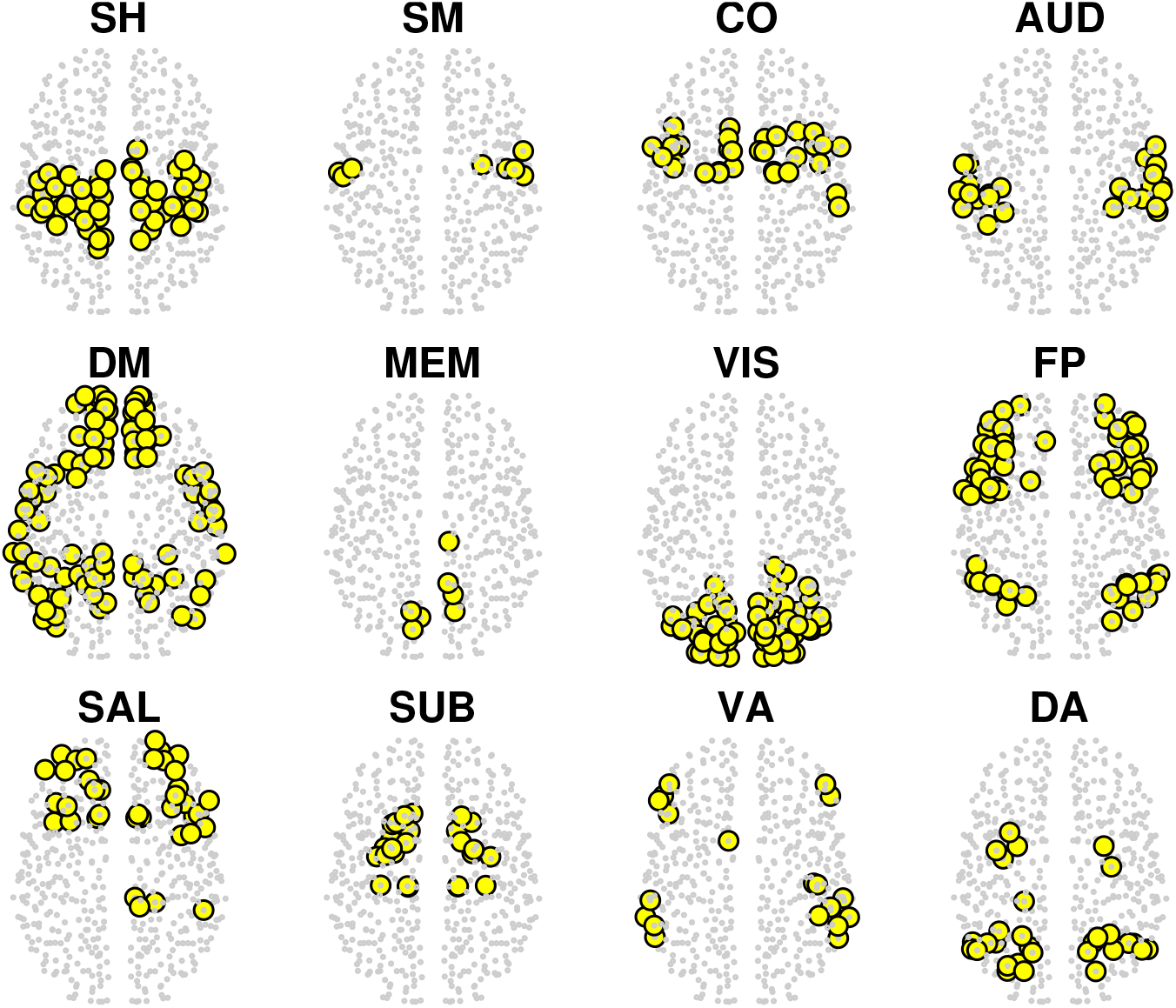
Spatial map of the connectome’s functional subnetworks (in superior axial view). SH=Somatomotor Hand, SM=Somatomotor Mouth, CO=Cingulo-Opercular, AUD=Auditory, DM=Default Mode, MEM=Memory, VIS=Visual, FP=Fronto-Parietal, SAL=Salience, SUB=Subcortical, VA=Ventral Attention, and DA=Dorsal Attention.

**Figure S14:**
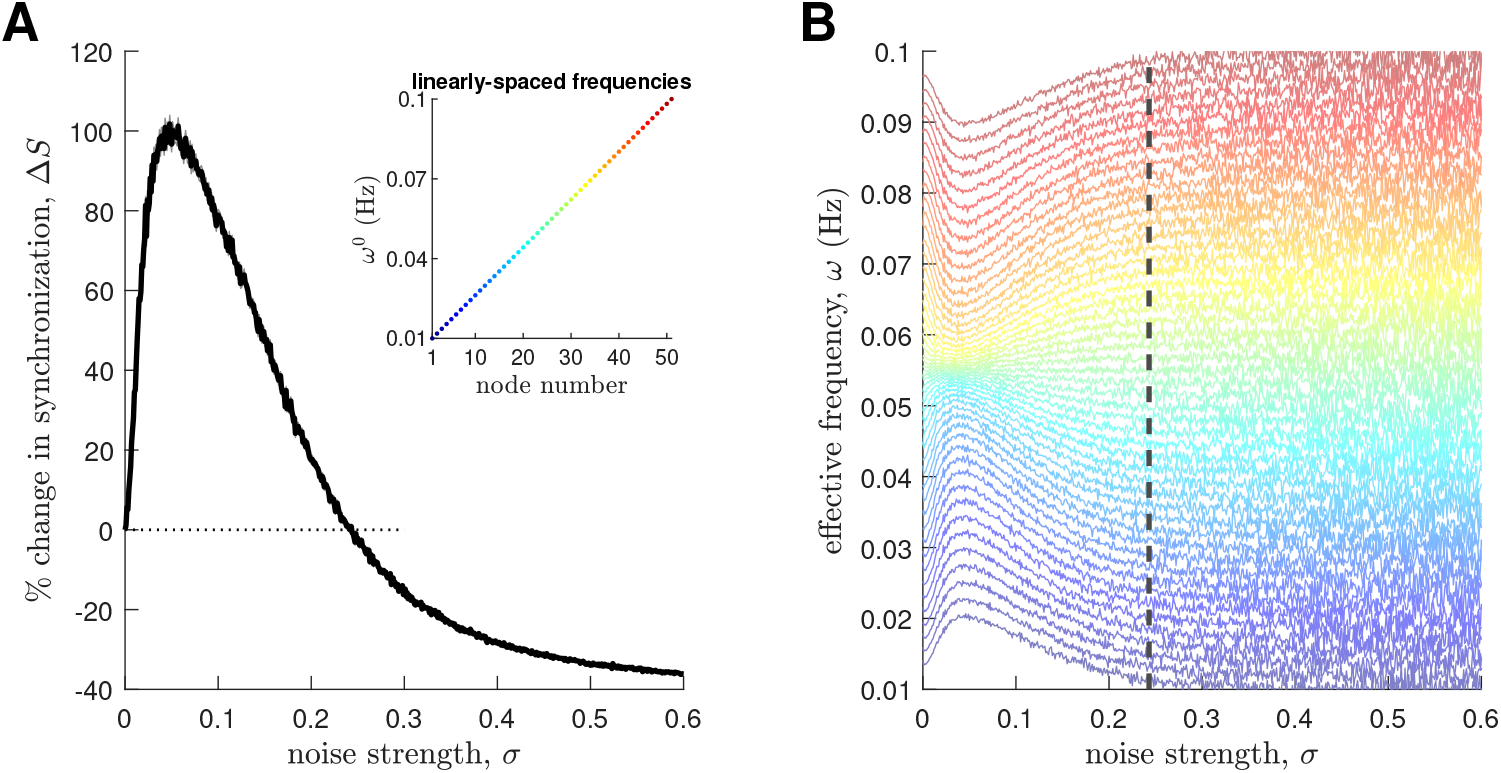
Synthetic model of a fully connected network (*N* = 51) with linearly-spaced natural frequencies from *ω*^0^ = 0.01 to 0.1 Hz. **(A)** Percent change of network synchronization Δ*S* vs noise strength *σ*. The solid line represents an ensemble average of 50 noise realizations and the shaded area represents the standard error of the mean. The inset shows the natural frequencies, where the markers are colored according to their order (blue: low *ω*^0^; red: high *ω*^0^). **(B)** Effective node frequency *ω* vs noise strength *σ*. The lines are colored similarly to the inset of panel A. The lines also represent ensemble averages of 50 noise realizations. The dashed line represents the *σ* where Δ*S* in panel A becomes negative.

**Figure S15:**
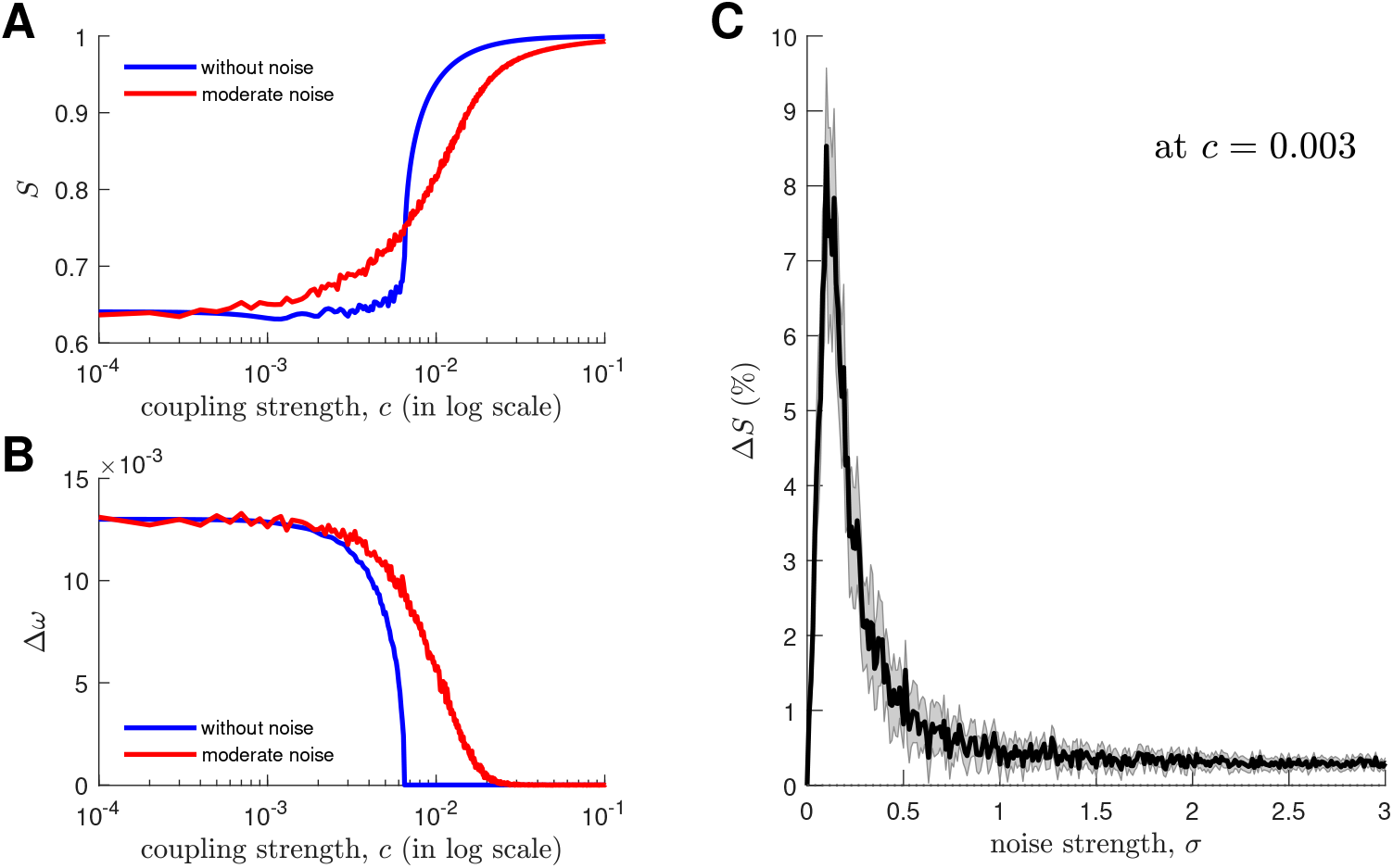
Synthetic model of two connected oscillators with natural frequencies of 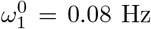 and 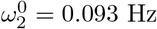. **(A)** Synchronization *S* vs coupling strength *c* for two noise strengths (blue: without noise; red: moderate noise). **(B)** Difference in effective node frequencies Δ*ω* (= *ω_i_* – *ω*_2_) for two noise strengths (blue: without noise; red: moderate noise). For panels A and B, the red solid line represents an ensemble average of 50 noise realizations. **(C)** Percent change of synchronization Δ*S* vs noise strength *σ* at *c* = 0.003. The solid line represents an ensemble average of 50 noise realizations and the shaded area represents the standard error of the mean.

**Figure S16:**
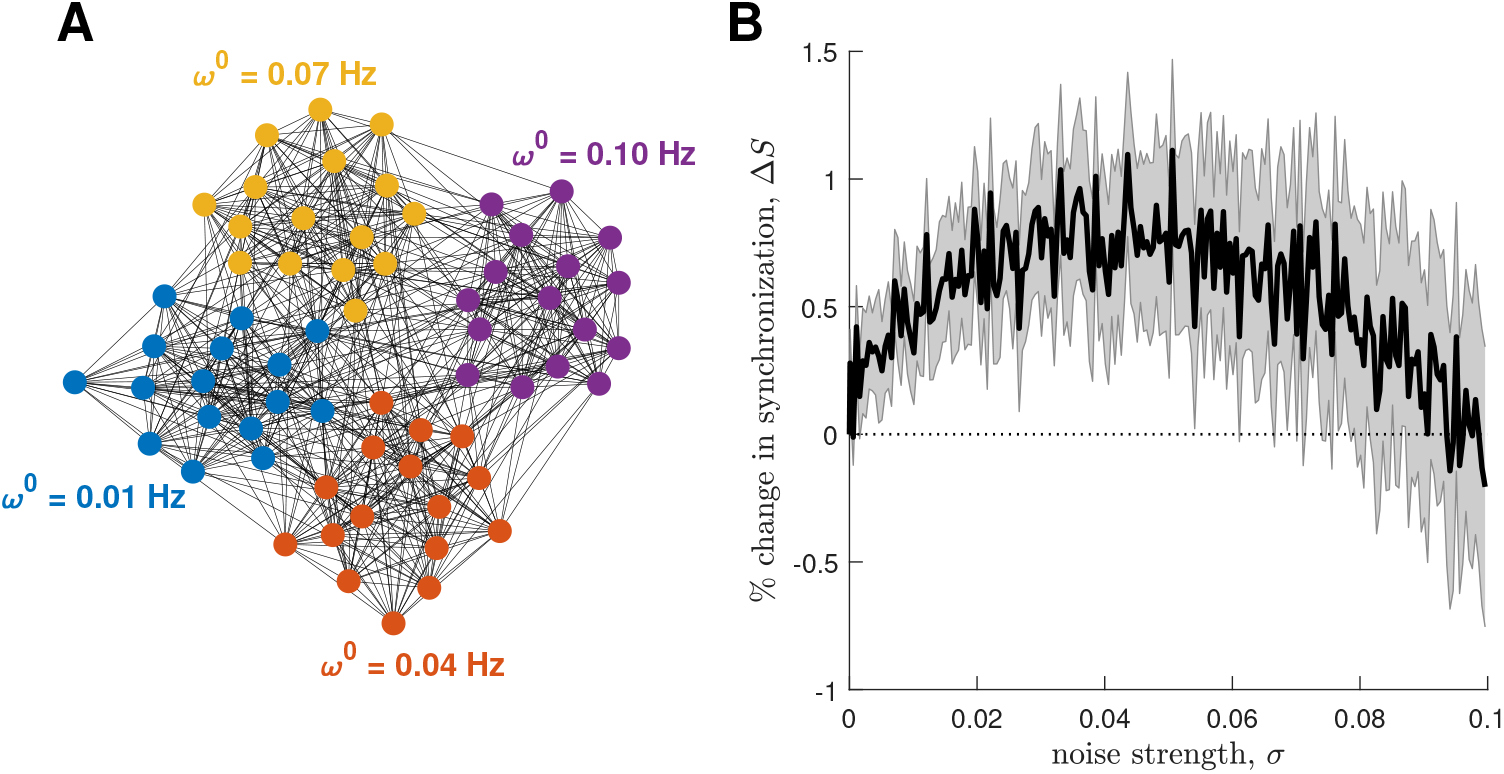
Synthetic model of a random modular network (*N* = 64). (A) Network with four modules. Within-module connectivity is all-to-all, while between-module edges are randomly created until the entire network has 1200 edges (network density ~ 30%). The colors represent nodes belonging to the same module. All nodes within each module oscillate at the same intrinsic natural frequency (i.e., *ω*^0^ = 0.01, 0.04, 0.07, 0.10 Hz for each of the modules). (B) Percent change of global network synchronization Δ*S* vs noise strength *σ*. The solid line represents an ensemble average of 50 network surrogates and the shaded area represents the standard error of the mean.

**Figure S17:**
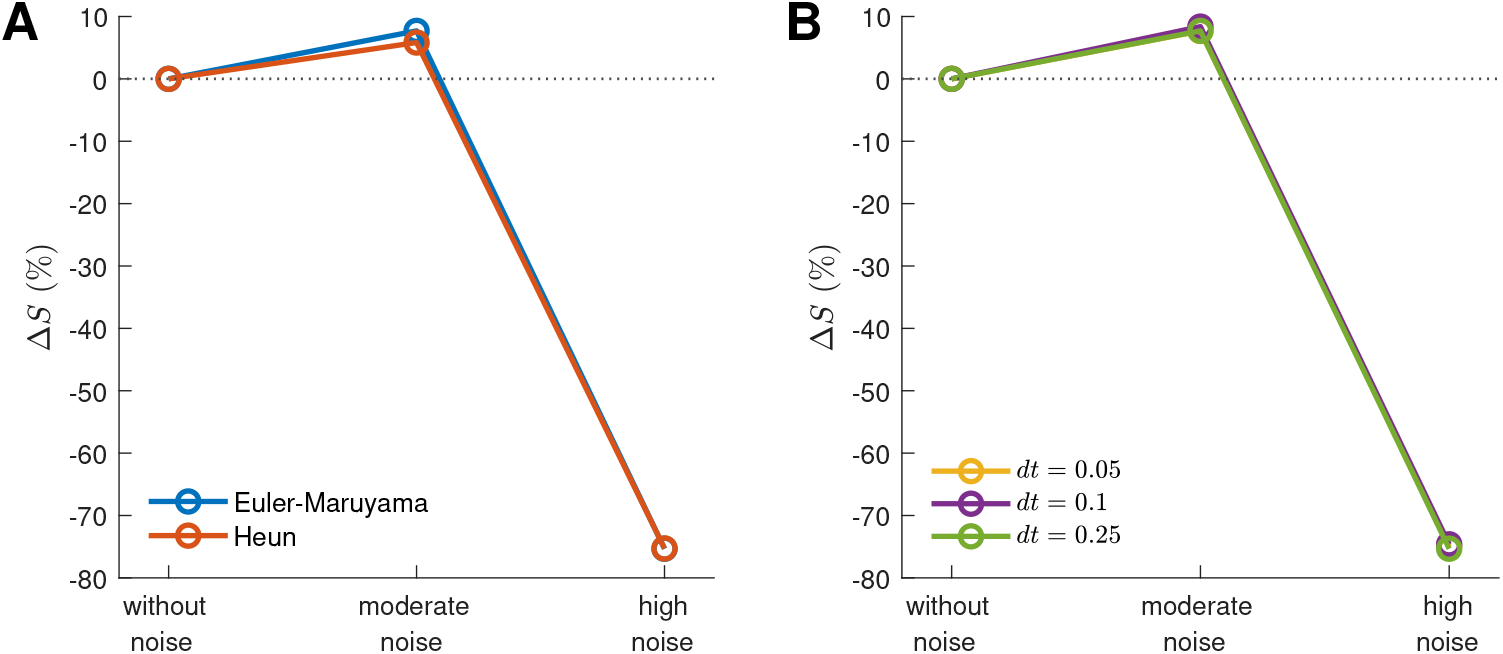
Robustness of the stochastic synchronization effect to the choice of **(A)** numerical integration scheme and **(B)** timestep *dt*. The markers represent ensemble averages of 50 noise realizations.

**Figure S18:**
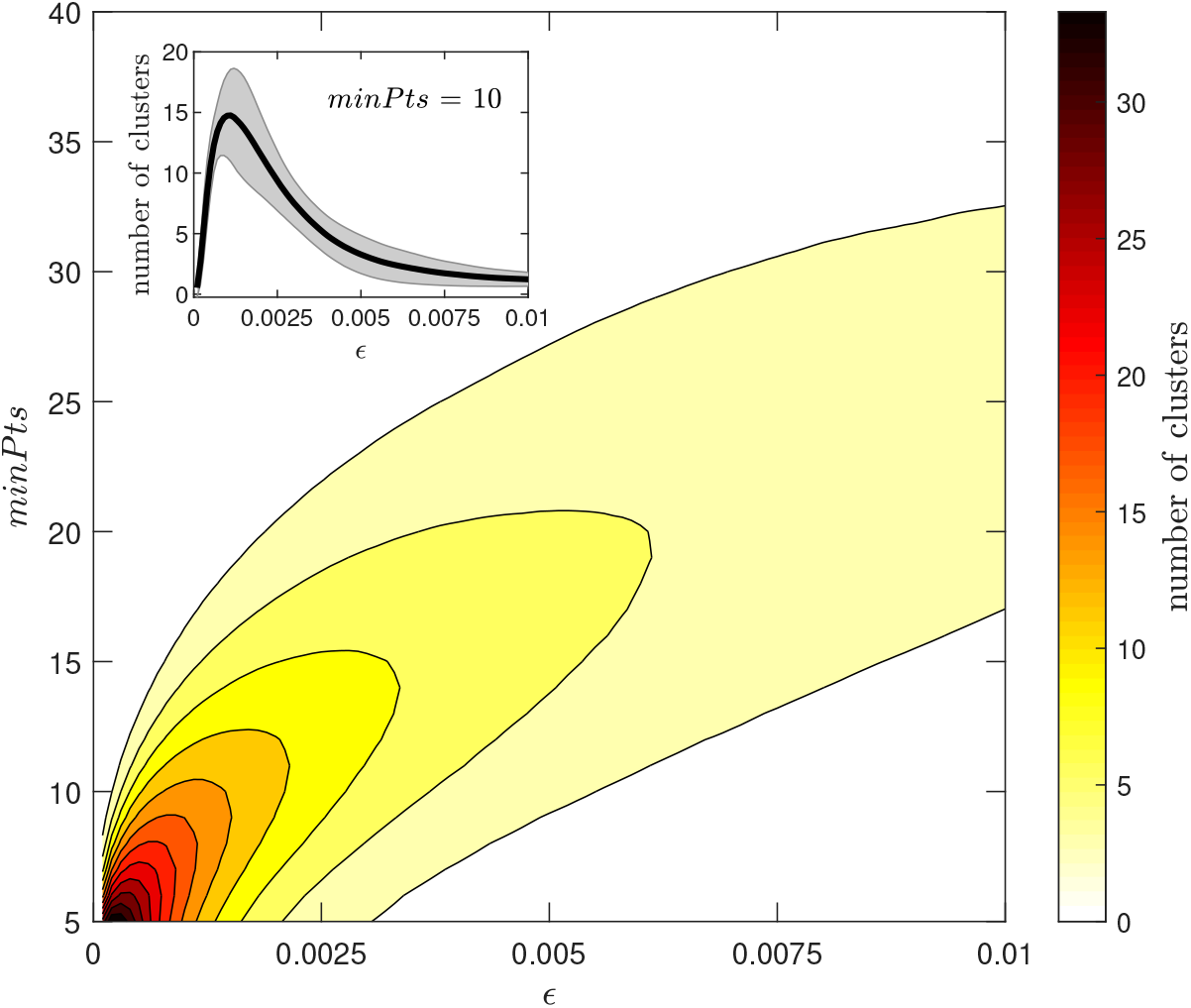
Optimization of the DBSCAN clustering algorithm. The number of clusters for the noise-free case is calculated for each combination of cluster radius e and minimum cluster size *minPts.* **(Inset)** Number of clusters vs *ϵ* at *minPts* = 10. The solid line represents the time average and the shaded area represents the standard deviation.

Movie S1: Network coherence and node phases for dynamics without noise. **(Top)** Time evolution of network coherence *R* from *t* = 9000–10000 s. **(Bottom)** Spatial distribution of node phases *θ* in different conventional brain views (left: superior axial; middle: sagittal; right: posterior coronal).

Movie S2: Network coherence and node phases for dynamics with moderate noise. **(Top)** Time evolution of network coherence *R* from *t* = 9000–10000 s. **(Bottom)** Spatial distribution of node phases *θ* in different conventional brain views (left: superior axial; middle: sagittal; right: posterior coronal).

Movie S3: Network coherence and node phases for dynamics with high noise. **(Top)** Time evolution of network coherence *R* from *t* = 9000–10000 s. **(Bottom)** Spatial distribution of node phases *θ* in different conventional brain views (left: superior axial; middle: sagittal; right: posterior coronal).

